# A hawkmoth’s point of view - Innate symmetry preferences are informed by floral display and foraging function in an insect pollinator

**DOI:** 10.64898/2026.11.04.764239

**Authors:** Anupama Nayak Manel, Lochlan Walsh, Anna Stöckl

## Abstract

Floral visual cues are key for flower recognition and feeding in many pollinators. In insects, innate preferences for characteristic floral colour and pattern cues can guide naïve foragers during their initial foraging. Symmetry is suggested to be one such foraging cue, since it is a remarkably conserved floral feature, with flowers mainly displaying radial or bilateral morphologies. Insect pollinators like honeybees, bumblebees, and hoverflies have been found to distinguish symmetry from asymmetry spontaneously or through learning. However, whether innate preferences for floral symmetry types, i.e. radial or bilateral, influence naïve flower choice, and whether such preferences are linked to functional benefits during foraging remains uncertain.

We investigated innate symmetry choice and its associated foraging function in the hummingbird hawkmoth (*Macroglossum stellatarum*). Besides relying on pattern cues for flower choice, these hawkmoths feed while hovering, making symmetry a valuable visual cue for flight and proboscis control. In choice assays, the hawkmoths’ innate preferences for symmetry types depended on the floral display planes, with considerable differences between the horizontal and vertical displays. Tracking the hawkmoths during foraging revealed that in both display planes, the innately preferred patterns increased foraging success by guiding approach and nectary search. While an analysis of flowers visited by hawkmoths in nature did not reveal the same link between symmetry and display plane, a majority of plant species with vertically displayed flowers showed features matching those innately preferred by the hawkmoths. In this study, we provide the first instance of display-dependent innate foraging preferences and highlight how visuomotor strategies in nectivorous insects may have coevolved with flower features.

## Introduction

Insect pollinators must detect food sources in highly cluttered environments, for which they use flower-specific odour, colour and pattern cues. Amongst these floral cues, flower patterns may play a crucial role in guiding flower choice for naïve foragers with no prior experience of flowers. In several pollinator species such as honeybees (Lehrer et al., 1997), bumblebees (Lunau, 1991; Simonds & Plowright, 2004), hoverflies (Mishra et al., 2025; Møller & Sorci, 1998), and hawkmoths (Kelber, 1997, 2002), spontaneous pattern preferences have been demonstrated in the context of foraging, where, with no prior experience of the patterns, there exist biases in choice towards particular pattern types. These spontaneously preferred patterns reflect visual characteristics found in flowers, such as radial spoked patterns (Kelber, 1997; Lehrer et al., 1997; Simonds & Plowright, 2004) or yellow central elements (Dinkel & Lunau, 2001; Kelber, 1997) suggesting that spontaneous preferences for ‘flower-like’ patterns play a key role in guiding inexperienced pollinators to identify their initial food sources.

These innate pattern preferences are thought to be the result of the co-evolution of flowering plants and insect pollinators, with both sides mutually benefiting during foraging interactions. For flower foragers, functional benefits from floral patterns might occur at various phases of foraging, from aiding flower selection and guiding the forager’s approach and landing (Lunau, 1990; Richter et al., 2023; Wiegel & Lunau, 2026) to enabling faster nectary or pollen discovery (Dinkel & Lunau, 2001; Goyret, 2010; Johnson & Dafni, 1998; Lawson et al., 2017; Leonard & Papaj, 2011). For plants, displaying specific patterns can improve pollination success by guiding the foragers’ movements to increase the probability that they contact the sexual organs of the plant (Hansen et al., 2011; Hempel de Ibarra et al., 2015; Wiegel & Lunau, 2026). Additionally, flowers with patterns were shown to have greater flower fidelity, with higher visitation rates even when reward quality was lower (Leonard & Papaj, 2011).

While spontaneous pattern preferences and the influence of patterns on foraging have been studied separately, there is no evidence to show that innately preferred patterns offer additional foraging function. A visual feature which is a likely candidate to study this link between innate flower pattern choice and foraging function is floral symmetry. A majority of angiosperm flowers are symmetric, with flower morphologies and their associated patterns being either bilaterally or radially symmetric (Neal et al., 1998). Whether a flower is radially or bilaterally symmetric is linked to the plane in which the flowers are displayed, such that plant species with vertically displayed flowers typically show bilateral symmetry while horizontally displayed flowers typically show radial symmetry (Fenster et al., 2009; Neal et al., 1998). Given the ancestral trait of flowers being radially symmetric (Sauquet et al., 2017), the emergence of bilaterally symmetric flowers is speculated to be linked to pollinator-mediated selection. Such pollinator selection could emerge from the influence of symmetry on foraging efficiency. In honeybees, symmetry type was found to influence handling time, with less time spent on artificial bilateral flowers compared to radial flowers (West & Laverty, 1998). Symmetry may also signal reward, since in some plant species, the available nectar volume is linked to the degree of floral symmetry (Møller, 1995; Møller & Eriksson, 1995). The ability to distinguish symmetric from asymmetric features during foraging has been shown in pollinators such as the honeybee (Giurfa et al., 1996; Wignall et al., 2006), bumblebee (Hempel de Ibarra et al., 2015; Plowright et al., 2017), and hoverfly (Mishra et al., 2025; Møller & Sorci, 1998). In nature, higher visitation rates for flowers with greater symmetry have also been recorded (Møller & Eriksson, 1995). However, whether pollinators innately have preferences for bilateral or radial symmetry types remains uncertain.

To quantify innate symmetry preferences and their functions in a flower-foraging insect, we studied the hummingbird hawkmoth (*Macroglossum stellatarum*). From flower choice to feeding from the nectary, vision plays a key role in guiding foraging behaviour in this diurnal hawkmoth (Goyret & Kelber, 2012; Kannegieser et al., 2024; Kelber, 2002). Innate foraging choices could be elicited from these hawkmoths with no prior habituation or feeding experiences using artificial floral stimuli, where blue and yellow contrasting patterns with radial spokes were found to attract robust foraging choices (Kelber, 1997). Visual information also plays a key role in guiding the motor control of flower interactions in these hawkmoths, particularly to manoeuvre their long proboscis into the nectary while hovering in front of the flower (Goyret & Kelber, 2012; Kannegieser et al., 2024; Kelber, 2002).

In this study, we assessed innate preferences for floral symmetry types to then test the influence of preferred patterns on the hawkmoths’ foraging strategies and feeding efficiency across the vertical and horizontal floral display plane. We reconciled the results obtained from artificial flowers with an analysis of the symmetry types of flowers naturally visited by this hawkmoth species.

## Materials and Methods

### Animals

Hummingbird hawkmoth (*Macroglossum stellatarum*) populations were raised indoors at 23 °C and 40-60% humidity with a 14:10 h light:dark cycle. The hawkmoth populations were maintained at the University of Konstanz, Germany since 2022 and supplemented in summer each year with up to a dozen moths captured in Konstanz. Eggs were collected from the lab populations and wild-caught females when available. Caterpillars were raised on their native host plant *Gallium* sp.. Adult moths were fed in groups in feeding cages (60 cm × 60 cm × 60 cm) with 20% (w/v) sucrose-water solution from gravity feeders.

### Experimental Setup

Experiments were performed in a flight cage of 60 × 60 × 60 cm dimension at 23 °C. The cage was lit from above with four 60 cm long daylight-like fluorescent tubes (Osram L 18 W/965 Biolux Tageslicht G13). Lights were connected to an electrical ballast (GloMat 2 × 40 W, Hagen) to increase the flicker frequency of the fluorescent tubes above 25 kHz, beyond the resolvable range of hawkmoths (Stöckl et al., 2019).

Artificial flower patterns in Blue (CMYK 91:66:0:0) and Yellow (CMYK 0:0:100:0) were designed in the vector graphics software Inkscape (Inkscape v1.3 2023, https://inkscape.org). The flower patterns were 38 mm in diameter with yellow features on a blue background. The selected colours were within the spectral range of hawkmoth colour vision (Telles et al., 2014) and the particular size and colours of the artificial flower patterns were chosen to elicit spontaneous and robust foraging approaches from the hawkmoths during experiments (Kelber, 1996, 2010). The colours were laser printed (imageRUNNER ADVANCE DX C3926i, Canon) on unbleached paper (‘Classic White’, Steinbeis) and laminated with matte foil (S-PP525-22 matt, Peach, PRT GmbH). To provide suitable optic flow during flight, a matte-laminated sheet with dead-leaves patterns (Lee et al., 2001) was provided in the cage. A dead-leaves pattern covered cardboard platform (30 × 50 × 50 cm) in the cage was used to decrease the flight height and thereby obtain rapid foraging responses.

### Display planes and pattern sets

Pattern sets were presented either in the horizontal or the vertical display plane to emulate two common floral display planes. In the horizontal display plane, artificial flowers were constructed by pasting the pattern discs on cylindrical plastic pedestals (8 x 3.5 cm) wrapped with unbleached paper. The flowers were placed on a cardboard platform (30 × 50 × 50 cm) covered with the dead-leaves patterns (Fig. 1a). In the vertical display plane, patterns were fixed on the dead-leaves background across two adjacent vertical walls of the flight cage. Since moths tended to choose the pattern closest to the roof of the flight cage, all patterns were presented on the same level across a row (Fig. 1a). Each of the cage walls was lined with one of each pattern type from the pattern set, and while repositioning, the pattern height from the cage floor was maintained.

**Fig. 1.**
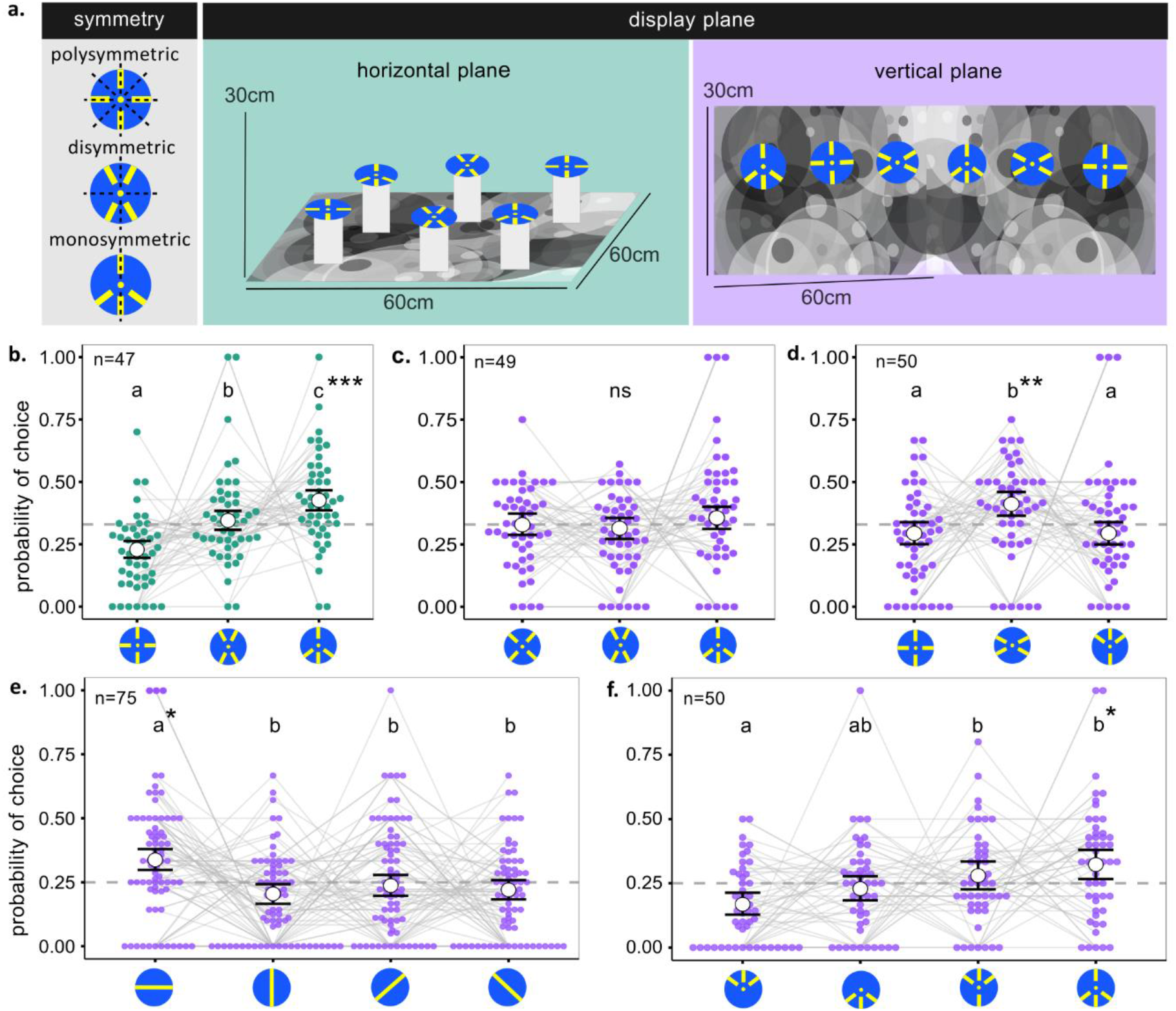
Naïve floral pattern choice. **a.** Patterns with variations in symmetry presented in the horizontal (green) and vertical (purple) display planes. **b-f.** Probability of naïve pattern choices by hawkmoths are represented by white circles with error bars representing 95% Credible Intervals. Coloured points represent the proportion of choice from individual hawkmoths, with the grey lines connecting choice proportions across the same individual. Naïve foraging preferences were tested across pattern symmetries in the horizontal display plane (b), and in the vertical display plane across 2 sets with varied orientations (c,d). Naïve foraging preferences in the vertical display plane were tested across bar orientations (e) and feature positions (f). The number of hawkmoths tested in each pattern set is given by ‘n’. Letters represent significant differences between pattern types (Table 1), and the patterns chosen significantly above random chance are denoted by asterisks *p < 0.05; **p < 0.01; ***p < 0.001.

**Fig. 2.**
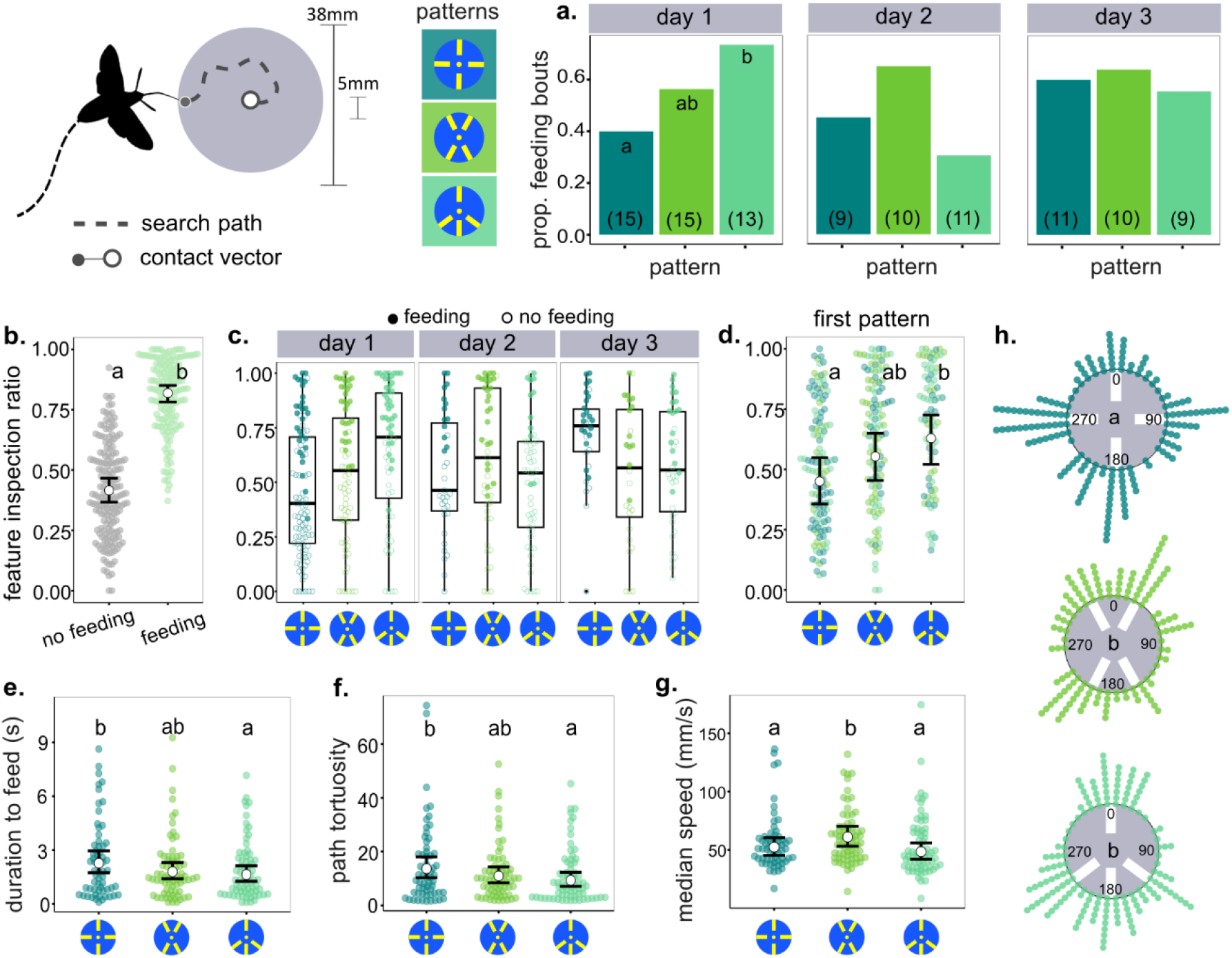
Foraging interactions with horizontally displayed symmetry patterns. **a.** Proportion of pattern interactions with successful feeding across symmetry types on each day of testing. The values at the x-axis indicate the number of hawkmoths tested. **b-d.** Ratio of the proboscis contacts on the yellow bar features compared with the total proboscis contacts on the flower during inspection. The feature inspection ratio is compared across bouts with and without feeding (b.), the respective symmetry types on each day of testing (c.), and the symmetry type that was interacted with on day 1 (d.). The predicted means from the GLMMs with significant differences are represented by the white circle, and error bars indicate 95% Confidence Intervals. Groups with no differences (p> 0.05) share the same letter. **e-g.** Proboscis path dynamics across symmetry types from bouts with feeding across all days. Distribution of time taken to feed from the central nectary (e.), tortuosity of the proboscis path (f.), and mean proboscis speed (g.), with predicted means from the GLMM denoted by white circles and the error bars depicting 95% Confidence Intervals. Pattern types with no differences (p> 0.05) share the same letter. h. Distribution of first proboscis contacts with respect to the pattern centre. Contact vector distributions with no differences as indicated by a Watson’s 2-sample test (p> 0.05) share the same letter.

**Fig. 3.**
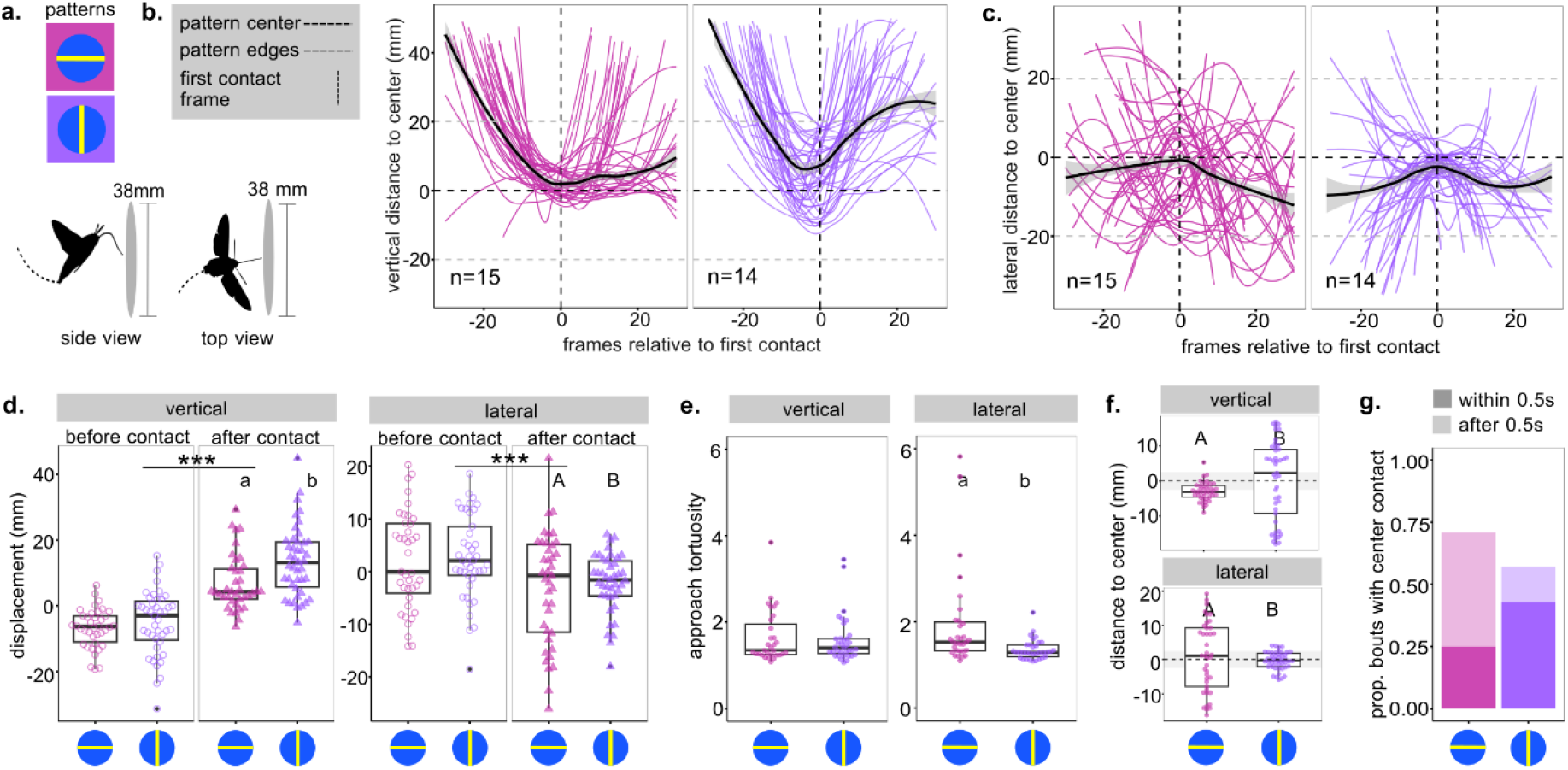
Foraging interactions with vertically displayed patterns across two bar orientations. **a.** The bar orientation patterns were filmed from the side to record the vertical movement, and from the top to record lateral movement. **b-c.** Distance of the hawkmoth from the centre of the pattern across 30 frames (1 s) before and after the first contact with the pattern in the vertical (a.) and lateral (b.) plane. Coloured lines depict smoothened individual trajectories. The black line represents the smoothened median coordinate values, with the grey shaded regions denoting the standard error. The number of individuals interacting with each pattern type is given by ‘n’. **d.** Displacement of the moth 0.5 s before and after first proboscis contact with the pattern. Positive values indicate upward movement, and negative values indicate downward movement in the vertical plane and leftward and rightward movement, respectively, in the lateral plane. Asterisks denote significant differences between the groups (p<0.001). Lowercase letters denote significant differences (p<0.05) in means from the GLMM, and uppercase letters denote significant differences in distribution of the data from the Levene’s test. **e.** Tortuosity of the approach trajectories up to the first pattern contact across pattern types. Letters denote significant differences in means (p<0.05) from the GLMM **f.** Vertical distance from the pattern centre of the moths’ proboscis tip during the first pattern contact. The grey shaded region depicts the central circle on the pattern representing the floral nectary. Predicted means from the GLMM are represented by the white circle, and error bars indicate 95% Confidence Intervals. Letters denote significant differences (p<0.05) in the distribution of the data from the Levene’s test. **g.** Proportion of bouts where the proboscis contacted the pattern centre. The darker shades represent the inspections that contacted the centre within 0.5 s of the first contact.

**Table 1.** Pairwise naïve pattern choice comparison between patterns in each pattern set. Significant pairwise differences are highlighted in bold.

| **Display position** | **Pattern set** | **Pairwise contrasts** | **Odds ratio** | **Estimate [95% Credible Interval]** | **P value** |
|---|---|---|---|---|---|
| horizontal | symmetry | polysymmetric : disymmetric | 0.66 | -0.41 [-0.64, -0.19] | **>0.001** |
|  |  | polysymmetric : monosymmetric | 0.53 | -0.62 [-0.85, -0.41] | **>0.001** |
|  |  | dissymmetric .: monosymmetric | 0.81 | -0.21 [-0.40, -0.03] | 0.024 |
| vertical | symmetry set 1 | polysymmetric : disymmetric | 1.04 | 0.04 [-0.18, 0.27] | 0.688 |
|  |  | polysymmetric : monosymmetric | 0.92 | -0.08 [-0.30, 0.15] | 0.688 |
|  |  | dissymmetric .: monosymmetric | 0.88 | -0.13 [-0.36, 0.10] | 0.688 |
|  | symmetry set 2 | polysymmetric : disymmetric | 0.71 | -0.33 [-0.58, -0.11] | **0.008** |
|  |  | polysymmetric : monosymmetric | 1.00 | 0 [-0.26, 0.26] | 0.999 |
|  |  | dissymmetric .: monosymmetric | 1.40 | 0.34 [0.10, 0.57] | **0.008** |
|  | bar orientation | 0° : 90° | 1.64 | 0.49 [0.24, 0.76] | **0.001** |
|  |  | 0° : 45° | 1.42 | 0.35 [0.09, 0.61] | **0.015** |
|  |  | 0° : -45° | 1.52 | 0.42 [0.19, 0.67] | **0.002** |
|  |  | 90° : 45° | 0.86 | -0.14 [-0.45, 0.15] | 0.533 |
|  |  | 90° : -45° | 0.93 | -0.07 [-0.36, 0.21] | 0.638 |
|  |  | 45°: -45° | 1.10 | 0.07 [-0.21, 0.36] | 0.638 |
|  | feature position | only top : only bottom | 0.74 | -0.30 [-0.68, 0.05] | 0.166 |
|  |  | only top : greater top | 0.60 | -0.51 [-0.86, -0.16] | **0.016** |
|  |  | only top : greater bottom | 0.52 | -0.65 [-0.21, 0.36] | **0.002** |
|  |  | only bottom : greater top | 0.81 | -0.20 [-0.53, 0.12] | 0.270 |
|  |  | only bottom : greater bottom | 0.71 | -0.34 [-0.69, -0.01] | 0.081 |
|  |  | greater top : greater bottom | 0.87 | -0.14 [-0.47, 0.17] | 0.411 |

Multiple pattern sets were designed, with each set testing foraging preferences between three or four patterns varying in specific pattern qualities. The patterns were designed as variations of the blue-yellow patterns previously found to attract strong foraging responses in naïve hawkmoths (Kelber et al. 1997), with the features based on the most attractive patterns found in pilot experiments (see supplementary Fig. S1). To test symmetry preferences, a pattern set was created with the yellow features repositioned to obtain three symmetry types– polysymmetry, dissymmetry, and monosymmetry (Fig. 1a), based on their planes of symmetry (Neal et al. 1998). The pattern features consisted of a central circle (2 mm radius) surrounded by rectangular arms (12 x 4 mm) (Fig. 1b-d), with feature dimensions that could be resolved by the hawkmoths (Stöckl et al., 2017). In the vertical display plane, the symmetry patterns were presented across two sets within which the orientations of the features were maintained (Fig. 1c-d).

To test pattern feature orientation preferences, 4mm thick yellow bars on the blue discs were displayed in the vertical plane 0°, 90°, 45°, and 135° (Fig. 1e). For the feature-position preference test, the monosymmetric patterns were manipulated to have the rectangular arm features only in the upper or lower half, or differ relatively in the number of features in the upper and lower half (Fig. 1f). Additional pattern sets tested preferences between dissymmetric cross orientations, and between the multi-and dissymmetric patterns at two orientations each, to test interactions between symmetry and orientation (Fig. S2 j, k).

### Experimental procedure

#### Naïve pattern choice

Feeding-naïve adult hawkmoths were collected within 12 hours after eclosion, during which they had sufficient time to dry their wings and perform their first flight bouts. The moths were placed under dark conditions in individual plastic vials for 24 hours to motivate foraging during the experiments (Kelber, 1966; Manel et al. 2026). No food was present during the pattern choice experiments, and after the experiments, the moths were moved to the feeding cages.

In a trial, a single naïve moth was placed at one end of the flight cage within a 10 cm wide strip along the cage wall, allowing at least 5 cm between the moth and the closest flower. The trial commenced when the moth began its flight, and choices were scored for 5 minutes of flight. The duration was limited to allow the moth to make multiple choices while controlling for the effect of unrewarded interactions on their choices. In every trial, moths were presented with two replicates of each pattern type within a pattern set. Each moth was tested for pattern choice within a single pattern set. The artificial flowers were arranged in a randomised order, spaced apart with at least 5 cm between the discs, and were repositioned across and between trials to minimise the influence of location cues on choices. A pattern choice was annotated for every separate approach to an artificial flower in which the proboscis contacted the pattern surface. Moths could make multiple choices during the trial, and trials with no choices within the first 3 minutes were excluded from the analysis. Sex of the moth and the time of start of a trial were noted. The cage floor and the pattern surfaces were cleaned periodically. A USB camera fixed to the upper corner of the cage was controlled by the ContaCam software (ContaCam v9.0.9beta4, https://www.contaware.com) to record the experiments at 30 fps to validate the manual scoring.

#### Horizontal display tracking

To assess whether the pattern type influenced approach and foraging success on horizontally displayed flowers, naïve moths were filmed while interacting with rewarded patterns from the symmetry pattern set. A single 12 mL drop of 20% (w/v) sugar was available at the centre of the pattern on the central yellow circle, which spatially represented the flower’s nectary. This drop was manually replenished after each feeding so that a sugar drop was always available during each separate pattern approach. A trial was recorded for 10 minutes or up to 4 feeding bouts, in which the moth drank the sugar droplet on the pattern. In every trial, a single pattern was presented 25 cm off the ground on a pedestal wrapped in white felt, and interactions were recorded with one naïve moth present in the cage. The patterns were always located at the centre of the cage and were rotated in place between individuals to avoid potential cage-location preferences affecting the approach analysis. Each moth was presented with a different symmetry type across 3 days such that all pattern presentation orders were similarly sampled across the tested group. Between each trial, the hawkmoth was placed under dark conditions in a marked vial. Hawkmoths that did not feed during the trial were manually fed on a neutral grey surface before being placed under dark conditions. The pattern interactions were filmed with a camera (acA1300-200uc, Basler) mounted directly above the pattern over the transparent cage roof (Kannegieser et al., 2024). The camera was focused on the pattern and controlled using the Pylon Viewer Software (Basler) at 200 frames per second (played back at 30fps), with a field of view of 496 x 500 pixels (ca. 12 x 12 cm), recording in grayscale (Mono 8 format). The recording was controlled manually by the experimenter to start when the hawkmoth began its approach to the pattern and to end when the individual left the pattern vicinity.

The videos were analysed with a trained DeepLabCut (ver. 3.0.2) network to automatically track the head, thorax, and proboscis tip of the hawkmoth (Walsh et al., 2026). The tracked videos were then manually validated using the MATLAB application DLTdv8 (Hendrick, 2008). A MATLAB script was used to rotate all the tracked coordinates to obtain the same reference frame for each pattern. Each trial was split into bouts for every separate approach to a pattern by splitting the video frames when the thorax coordinates jumped by over 50 pixels or were separated by over 10 untracked frames.

#### Vertical display tracking

To quantify the influence of pattern type when foraging from vertically displayed flowers, hawkmoths were presented with the patterns from the feature-position set, or with the 0° and 90° bar patterns from the bar-orientation set (see display planes and pattern sets), where naïve choices significantly differed. A single pattern was mounted on a wooden block (28 x 28 x 28 mm) positioned at the centre of one wall of the cage, which was covered with dead-leaves patterns. A USB camera with a standard lens fixed on one side of the cage and facing the pattern block, so that the pattern edge created a single vertical line in the camera view, was used to record the side view and capture movement in the vertical plane (Fig. 4a). Another camera mounted directly above the pattern was used to obtain a top view to record lateral movement in the trials with the bar-orientation set (Fig. 4a). In the side view, the camera covered a 12 cm horizontal space in front of the pattern and 4 cm above and below the pattern edges for the bar orientation set, and 16.8 cm horizontal distance and 6.5 cm space above and below the pattern in the pattern feature position set. In the lateral view, 11 cm in front of the pattern and 5 cm on either side of the pattern edges were recorded. The cameras were controlled by the Contacam software to record videos at 30 fps.

**Fig. 4.**
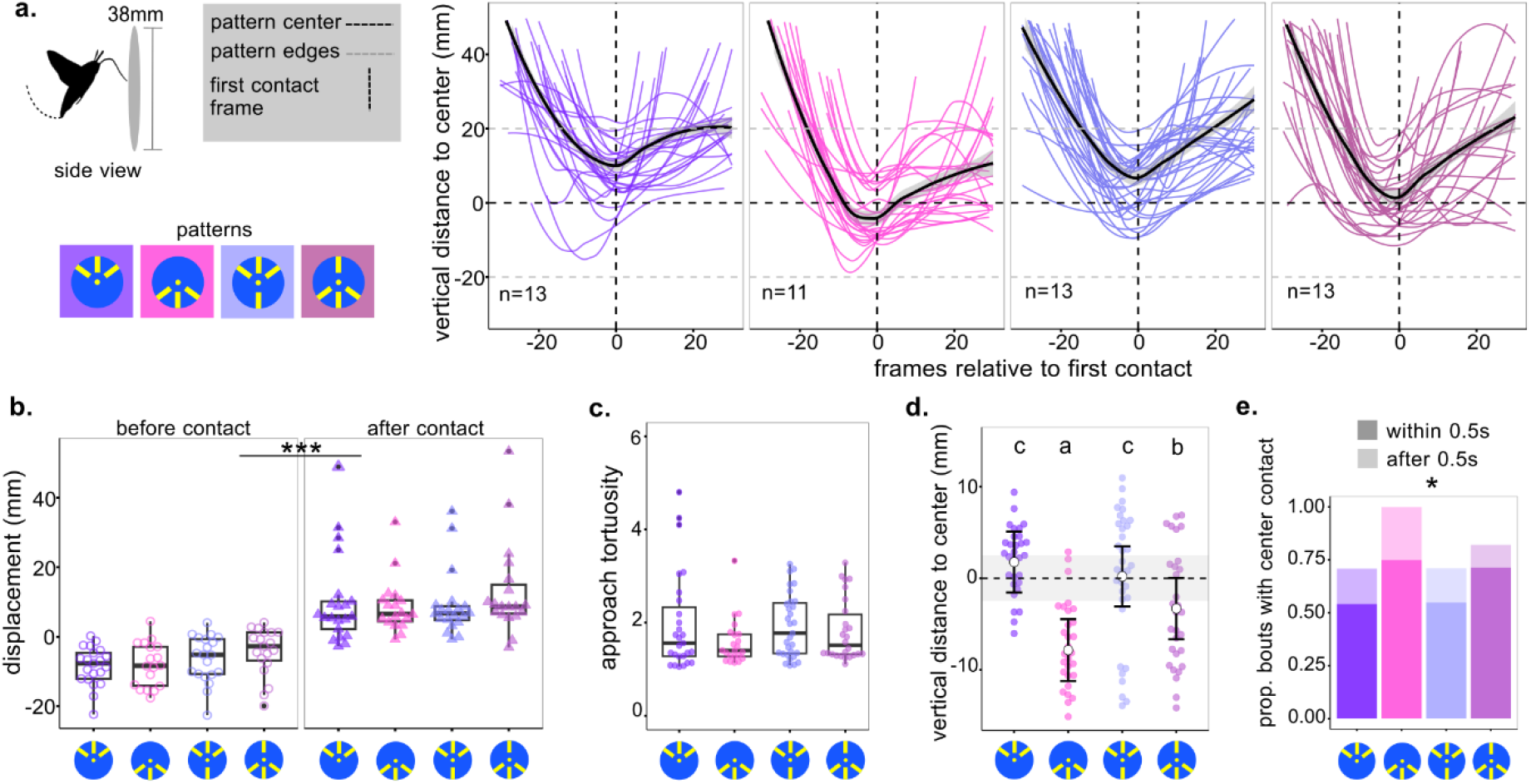
Foraging interactions with vertically displayed patterns across different feature positions **a.** Vertical distance of the moth’s body position from the centre of the pattern across 30 frames (1 s) before and after the first contact with the pattern. Coloured lines depict smoothed individual trajectories. The black line represents the smoothed median coordinate values, with the grey shaded regions denoting the standard error. The number of individuals interacting with each pattern type is given by ‘n’. **b.** Displacement of the moth 0.5 s before and after first proboscis contact with the pattern. Positive values indicate upward movement, and negative values indicate downward movement. Asterisks denote significant differences between the groups (p<0.001). **c.** Tortuosity of the approach trajectories up to the first pattern contact across pattern types. **d.** Vertical distance from the pattern centre of the moths’ proboscis tip during the first pattern contact. The grey shaded region depicts the central circle on the pattern representing the floral nectary. Predicted means from the GLMM are represented by the white circle, and error bars indicate 95% Confidence Intervals. Groups with no differences (p> 0.05) share the same letter. **e.** Proportion of bouts where the proboscis contacted the pattern centre. The darker shades represent the inspections that contacted the centre within 0.5 s of the first contact. Significant results from the Chi-square test based on the total proportion of bouts are denoted by an asterisk.

In a trial, the hawkmoth was placed in a random location 40cm from the cage wall with the pattern. A trial commenced when the hawkmoth started its flight and lasted for 3 min or up to 5 separate flight bouts in which the pattern was contacted by the proboscis. The video was then stopped, and the pattern was replaced with a different pattern from the set in a randomised order without disturbing the moth’s flight. Individual moths were tested with each pattern type within a single pattern set. Since it was not possible to have food rewards available in the vertical display without creating additional mechanical cues, these trials were unrewarded.

The videos were run through a custom-written MATLAB script to track the position of the moth using pixel differences between frames, from which the approach trajectory was estimated. The coordinates of the proboscis tip and head upon first contact with the pattern in every approach were manually annotated using the dtlDV8 application. Proboscis contacts after the initial contact were manually annotated in frames where the proboscis tip position on the pattern was clearly observable, allowing the quantification of the general direction of proboscis movement on the pattern.

#### Quantifying visual traits of natural flowers

Plant species visited by *M. stellatarum* were analysed to compare the hawkmoths’ artificial flower pattern preferences with natural flower shapes and patterns. Photographs of hummingbird hawkmoths visiting flowers were collected from a citizen science database, Naturgucker (naturgucker.de), to which data was uploaded between 2003 to 2024 from across Germany. Plant species were subsequently identified via the Flora Incognita algorithms (floraincognita.com/, Mäder at al. 2021) and retained for analysis if the identification certainty scores exceeded 0.7. A total of 2204 photographs were used in the analysis, with the highest representation from 3 common garden plants-*Buddleja davidii* (31%), *Phlox paniculata* (0.09%), and *Lavandula angustifolia* (0.05%). All other plant species each represented less than 0.05% of the photos in the dataset (Fig. S6). Since the number of photos could represent observer biases rather than indicating the hawkmoths’ foraging preferences, here, we assessed the dataset at the plant-species level to describe the range of floral visual features visited by this species.

Flowers from each species were annotated for pattern and shape features with respect to the two-dimensional visual features of the flower face, including the visible perianth, stamen, and pistil. Visual features of single flowers were annotated in clustered inflorescences such as umbels and corymbs. Multiple pictures of the flower were obtained through online searches and were assessed manually. The flower display plane was marked ‘horizontal’ when the petals were parallel to the ground, ‘vertical’ if orthogonal to the ground, and ‘downward’ if the corolla faced downwards, based on the display of the majority of flowers (>70%) on the plant (see examples in Fig. 5a). The display was considered ‘multiple’ when a majority of flowers did not fit any single category. Flower symmetry was defined based on the nature of the repetition of the corolla in relation to the centre of the flower’s receptacle, as modified from the description of perianth symmetry by Neal et al. 1998. The symmetry was ‘radial’ if there were more than two planes of symmetry around the receptacle (corresponding to the polysymmetric pattern), ‘bilateral’ for corollas with mono-or disymmetric arrangements, and ‘filled’ for flowers with spherical or conical receptacles (e.g. Thistle flowers) where the 2D plane around the receptacle could not be precisely annotated. Pattern presence on the corolla was marked when there were visible contrasting features, such as differences in colour or a visually apparent darker shade, for example, as created by mechanical contours. For flowers with a vertical display, pattern position was marked ‘uniform’ when patterns were present across the corolla, and ‘top-half’ or ‘bottom-half’ when the features were localised. Flowers with a prominently larger lower surface were marked as possessing a ‘lower-lip’. Lower-lip structures in flowers have been studied in relation to foraging success in birds (Smith et al., 1996; Temeles & Rankin, 2000) and bumblebees (Castellanos et al., 2004; Zung et al., 2015) and are suggested to serve as landing guides or nectary indicators.

**Fig. 5.**
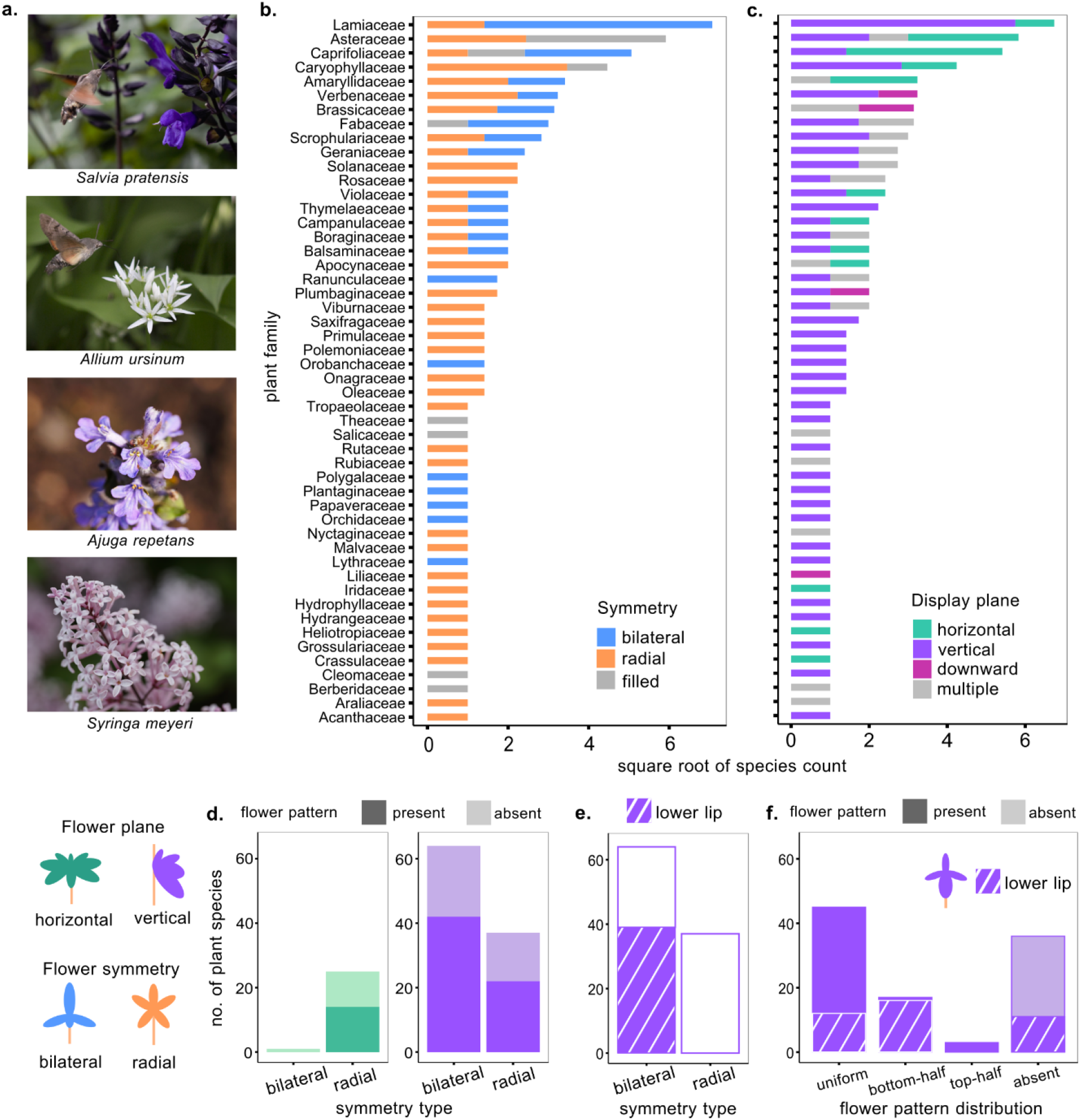
Characteristics of flowers foraged on by hummingbird hawkmoths. **a**. In order from top to bottom are examples of vertically displayed flowers, horizontally displayed flowers, bilaterally symmetric flowers with a lower lip, and radially symmetric flowers. **b-c**. Number of plant species with foraging visits across plant families. Colours depict symmetry type (b.) and display plane (c.) of the flowers. **d.** Number of plant species with bilateral and radial flower symmetries presented predominantly (>70%) in the horizontal display plane (left) and vertical display plane (right). **e.** Number of plant species within vertically displayed flowers with a lower lip. **f.** Number of plant species within vertically displayed flowers across pattern distribution types. Darker shaded regions represent the number of flowers with patterns, and the slanted white texture represents the number of flowers with a lower lip.

## Data Analysis

Statistical analyses were conducted in R (R Core Team, 2021), and plots were created with the ggplot2 package (Wickham, 2011).

### Pattern preference analysis

Probabilities of choosing a pattern type were calculated separately for each pattern set within its respective display plane. To account for the multinomial nature of the choice data, the proportion of pattern choices was modelled as multinomial GLMMs using a Bayesian framework. Sex was assigned as the predictor variable and animal ID as the random effect, and the models were fit with the priors set to the default distributions. The code was scripted with the Stan language (Carpenter et al., 2017; http://mc-stan.org/) via the brms package (v2.6.0; Bürkner, 2018; https://cran.r-project.org/web/packages/brms/) within R. The full model was compared using the leave-one-out (LOO) function (Vehtari et al., 2017) with a model where sex was dropped as a predictor. The model of best fit was assessed as the one with the highest estimated log predictive density (ELPD) value. The estimates were calculated from the final model assigned as

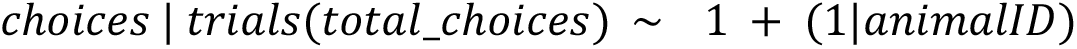

The parameters were estimated using four Markov–Chain Monte Carlo (MCMC) simulations from models where all chains converged (Rhat ≤ 1.01). Mean probability estimates and their credible intervals were calculated from the posterior draws of the fitted model coefficients taken from each iteration of the model and validated with the raw data means. The estimates were then compared to a null model assuming equal probabilities of choice across patterns (1/number of patterns) and adjusted for multiple testing with Benjamini-Hochberg corrections. To perform post hoc comparisons, we obtained pairwise contrasts by calculating the differences between the intercepts of every pattern pair at every draw on the log-odds scale. Z-scores (mean/SD) were drawn from these posterior distributions and analysed with a two-tailed test to derive p-values, which were adjusted using Benjamini-Hochberg corrections.

First choices were evaluated against a uniform probability distribution using a chi-square goodness of fit test. Variability in pattern selection across a sequence of choices within a trial was visualised across a rolling window of 3 choices for each animal to identify trends in selecting patterns across a trial period. The influence of a pattern choice on the next selection in the choice sequence was analysed using transition matrices.

### Horizontal display foraging interactions

MATLAB scripts were used to analyse the tracked videos and annotate successful bouts as approaches in which the moth’s proboscis arrived at the pattern centre, designated by a 2.5 mm radius circle, and remained within this centre for at least 20 consecutive frames, indicating feeding on the sugar droplet. Pattern feature inspection ratio in every bout was calculated as the proportion of proboscis contacts on the yellow bars with respect to the contacts on the background. The area of each bar was augmented by 2 mm on either side of the longer edges to include contacts on the feature edges as pattern feature inspections. Only contacts outside the central 2.5 mm circle region of the pattern were included in this ratio, and for successful bouts this was computed up to the moment of feeding to exclude post-feeding inspection.

In successful bouts, i.e., bouts with feeding, the proboscis tracks during inspection on the disc were used to calculate the duration required to arrive at the pattern centre after first pattern contact, and the median proboscis speed and path tortuosity up to feeding. Median approach angle with respect to the pattern centre was calculated from the approach flight tracks up to the first proboscis contact with the pattern. Only bouts where the moth was tracked from the boundaries of the recorded region, set to a distance of at least 40 mm from the pattern edge, were included in the approach trajectories, to exclude segments recorded after the animal was already close to the pattern. The position of the first contact on the pattern was quantified as the angle inscribed by the proboscis tip with respect to the pattern centre.

The data was analysed in R Studio. The influence of pattern type on bout success was analysed using a pairwise chi-square test of independence with Bonferroni corrections. Bout success was modelled as a response of feature inspection ratio to assess whether more precise feature inspection was linked to bout success. A generalised linear mixed model (GLMM) with the feature inspection ratio modelled as a betabinomial distribution was scripted using the ‘glmmTMB’ function. Bout success status was set as the predictor variable, and animal ID was set as the random effect to fit the model,

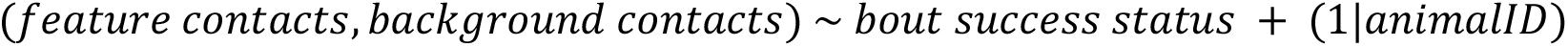

Dispersion parameters for the model were set to the bout success status to account for the heteroscedasticity in this predictor. Model fit was validated using the DHARMa package (Hartig, 2016). The ‘Anova’ function from the car package (Fox and & Weisberg, 2019) with Wald’s chi-square test was used to determine the significance of the predictor variable on the inspection ratios.

To compare pattern feature inspection ratios between pattern types, we used a GLMM with the feature contact ratios modelled as a betabinomial distribution. To account for pattern presentation order, the first presented pattern as a factor was set as one of the predictor variables. The model also contained the interaction between pattern type and trial day as a predictor variable and Animal ID as a random variable. Model fit was validated and stepwise reductions to the full model were performed with likelihood ratio tests (LRT) using the drop1 (Crawley, 2002) function. The simplest adequate model was selected as

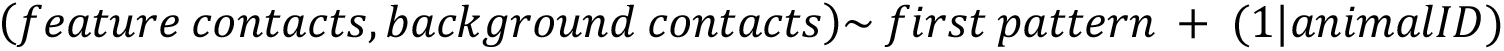

To check whether pattern inspection was asymmetrically directed in the monosymmetric pattern, the number of contacts on each rectangular arm in the pattern was compared using a chi-square test.

To assess how proboscis path dynamics of successful bouts varied with pattern type, we fit separate GLMMs with proboscis track duration and tortuosity as dependent variables. Approach durations across all bouts were also fit to a GLMM. In the models, the independent variables were assigned as the interaction between pattern type and trial day, and the first presented pattern as a factor to account for pattern presentation order. Animal ID was assigned as a random variable. The data was modelled as a Gamma distribution with a log link and fit with the ‘glmer’ function from the lme4 package. Model fit was validated and stepwise reductions to the full model were performed with likelihood ratio tests as with the previously described models. The simplest adequate models were selected as

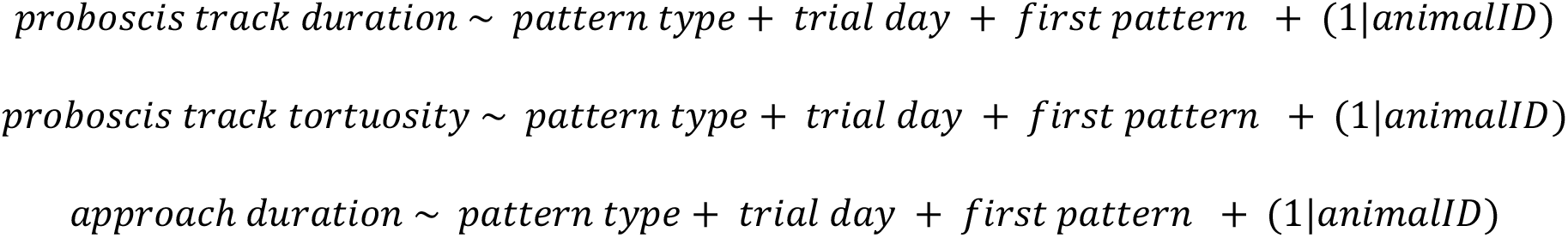

Similarly, an LMM was fit with log converted mean speed as the dependent variable, and was simplified using stepwise model reductions as

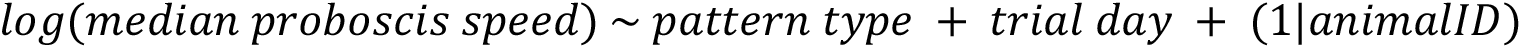

Post hoc comparisons from all models were drawn using the emmeans package and adjusted using Sidak corrections for multiple testing.

The approach and contact angle data was analysed using the ‘circular’ package in R. To test whether the approach and proboscis contacts were homogenously distributed around the patterns, a Rao’s spacing test was performed on the median approach angles and the proboscis contact angles using the ‘rao.spacing.test’ function. A Watson’s two-sample U^2^ test was performed to assess pairwise differences in angle distributions between pattern types using the ‘watson.two.test’ function.

### Vertical display foraging interactions

The data between pattern types were compared within their associated pattern sets, and values from the side and top view were also analysed separately. Trajectory coordinates relative to the pattern centre were calculated in MATLAB, and the relative y-coordinates from the tracks were used to visualise the movement trajectory across time (Fig. 4a,b). Since most tracked approaches were shorter than 1s (median [IQR]: 20 [17-34] frames), the approach trajectory metrics were visualised over a 20-frame period before the first pattern contact. The displacement across 0.5 s before the first contact was computed as the difference between the mean position of the hawkmoth across 15 frames before the first contact and its position at first contact. Displacement 0.5 s after first contact was given by the difference between the position at first contact and the mean position after 15 frames. The mean of the position was used to account for tracking errors. To compare displacement values before and after first contact across pattern types, a linear mixed model was fit with the ‘lmer’ function with the interaction between the pattern type and displacement instance (before/after contact) as the predictor, and animal ID as the random variable as,

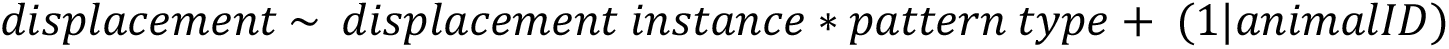

A Levene’s test was used to compare homogeneity of variances within the displacement data across pattern types within each displacement instance. Path tortuosity of the approach trajectories was calculated and compared between pattern types using a Wilcoxon’s test for the bar-orientation set with the 0° and 90° bars, and with a Kruskal-Wallis test for the patterns in the feature-position set. Only approaches with at least 5 tracked coordinates and with no greater than 4 continuous non-tracked coordinates were included in this analysis. Trajectory deviations at smaller temporal scales, i.e., path jitter, were estimated by Fourier transforming the approach coordinates to obtain power density distributions across frequencies. To compare jitter between the 0° and 90° bar patterns, where feature orientation was hypothesised to influence approach path stability, the power densities of frequencies lower than 5 Hz were assessed using a Wilcoxon’s test. However, with the temporal resolution of the video at 30 fps, trajectory frequencies lower than 15 Hz could not be computed in this analysis.

The manually annotated first proboscis contacts and the head positions relative to the pattern centre were compared across pattern types for differences in data variability using a Levene’s test. The position data was fit as a GLMM using the ‘glmmTMB’ function. The data for proboscis contact and head position were separately modelled as Gaussian distributions with the dispersion parameter set to the pattern types to account for data heteroscedasticity. The final comparisons were drawn from the model,

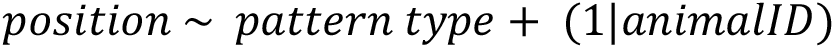

The occurrence of central nectary contact was assessed by checking whether, upon contact, the proboscis entered a 2.5 mm region from the pattern centre. This was quantified by computing the vectors between the first and last available annotated proboscis contacts across 10-frame windows within the whole proboscis inspection period in a bout. During inspection, moths would sometimes reinspect the pattern within the same bout upon reaching the upper edge of the disc by moving downwards and then upwards again (Fig.S4 c,d; S5 b). To quantify the duration to the first contact with the central nectary within a bout, the first proboscis contact with the central region was annotated. For the bar-orientation set, where both the side and top views of the proboscis inspection were available, a successful pattern centre contact was assigned only when the proboscis entered the pattern centre for corresponding bouts in both views. In the feature-position set, pattern centre contact was assessed only from the side view. A chi-square test was conducted on the total number of successful and unsuccessful bouts for each pattern type within a set to assess whether the pattern type influenced nectary discovery.

## Results

### Naïve pattern choices

Innate preferences for pattern symmetry type during foraging were recorded in hummingbird hawkmoths across two pattern display conditions, horizontal and vertical, to assess the influence of floral display planes on pattern choice. Naïve flower pattern choice with horizontally displayed artificial flowers was significantly influenced by symmetry type, with the monosymmetric pattern chosen above chance (posterior mean probability-0.43, 95%CrI [0.38 0.47], p<0.001; Table S1). The polysymmetric pattern was contacted significantly less frequently than both the monosymmetric pattern and the dissymmetric pattern (Table 1, Fig. 1b). This trend, however, was not replicated in the vertically displayed symmetry patterns where innate symmetry preference was tested with two sets of patterns, each with distinct and fixed orientations for every symmetry type (Fig.1 c-d) to account for the constrained pattern viewing angles. While there were no differences in choices between the symmetry types in the first set (Fig. 1c), within the second set, the dissymmetric pattern rotated to be parallel to the ground was chosen significantly above chance (posterior mean probability-0.42, 95%CrI [0.37 0.48], p=0.002; Table S1) and was contacted more often than both the mono-and polysymmetric patterns (Table 1, Fig.1d). There were no differences in choice probabilities between the polysymmetric and monosymmetric patterns in this set.

Pattern choices in the vertical display plane were thus not as clearly guided by symmetry as in the horizontal display plane but were instead influenced by other feature qualities. To identify the visual features that were relevant for choices in the vertical display plane, naïve moths were tested in four-way choice assays for preferences across upper vs lower feature positions, different bar orientations, various orientations of the dissymmetric pattern, and between polysymmetric and dissymmetric patterns in two different orientations. In the pattern set with different bar orientations, the bar oriented at 0° (parallel to the ground) was selected above chance (posterior mean probability-0.33, 95%CrI [0.26 0.40], p=0.036; Table S1), and with a greater probability than the other (90°, 45°,-45°) bar orientation patterns (Table 1, Fig. 1e).

There were no significant differences between the other presented bar orientations. In the upper vs lower feature position pattern set, the pattern with more features on the lower half was selected significantly above chance (posterior mean probability-0.32, 95%CrI [0.27 0.38], p=0.028; Table S1). Amongst the patterns within this set, the pattern with features only in the upper half was significantly less frequently selected than the patterns providing differential features in the upper and lower positions (Table 1, Fig. 1f). The other patterns were not chosen significantly differently from each other. In the remaining pattern sets that tested dissymmetric pattern orientations and interactions between symmetry and orientations, there were no biased pattern choices, and all patterns were selected at chance probabilities (Fig. S2j, k).

The first pattern choices from all the experiments were analysed to test whether performing a series of unrewarded pattern choices across 5 minutes led to extinction of the hawkmoths’ naïve preferences over time, resulting in differences between first choices and overall choices during the 5 minutes of testing. The distribution of first pattern choice in the horizontally displayed symmetry set (Fig. S2a) and one of the vertically displayed pattern sets (Fig. S2g) reflected the same trend as the analysis of pattern choice probabilities from all selected patterns. First choices from another of the vertically displayed sets varied from the complete pattern choice analysis and followed a similar trend to the choices made in the horizontal display (Fig. S2d). However, first choices were not significantly different between patterns in any of the symmetry pattern sets. First choices from the four-way assays in the vertical display plane also followed the same trend as the complete choice analysis, although the count differences between patterns were only significant for the bar orientation set (χ_3_² = 9, p =0.29, Fig. S2m).

Further, we assessed whether pattern choices across the 5 min duration changed consistently, indicating an extinction or change of preferences over consecutive non-rewarded choices. No consistent trend in the variability of pattern choices across the 5 min was found (Fig. S2b, e, h). The transition matrices for the pattern choices in the horizontal symmetry set (Fig. S2c) indicated a higher probability of choosing a preferred pattern after selecting an unpreferred pattern, and relatively lower probabilities of sequentially contacting the same pattern type. The transition probabilities between pattern types in the vertical display were random in the first set (Fig. S2f), while probabilities in the other set (Fig. S2i) indicated a high chance of sequentially contacting the dissymmetric pattern. We therefore consider the pattern choices during the entire 5 min test duration representative of the animal’s underlying preferences.

Thus, naïve hawkmoths preferentially selected monosymmetric patterns in the horizontal display plane, while in the vertical display plane, orientation and position of the pattern features, rather than the number of symmetry axes, were relevant for flower choice.

### Foraging efficiency on horizontally displayed pattern symmetries

To assess whether naïvely preferred floral symmetries offered functional benefits for foraging, feeding naïve hawkmoths were recorded interacting with horizontally displayed patterns across 3 symmetries – polysymmetry, dissymmetry, and monosymmetry. Animals interacted with a single unique pattern in a trial and encountered a different symmetry type across 3 days. The proportion of successful foraging bouts, i.e., bouts where the pattern inspections resulted in feeding from the central sugar droplet, was significantly influenced by the pattern type on the first day of pattern interaction (X ^2^= 13.53, p = 0.001) with more successful bouts on the monosymmetric pattern than on the polysymmetric pattern (p=0.002, Fig. 2a, day 1). There were no significant differences in proportion of successful bouts between symmetry types in the subsequent trial days (Fig. 2a).

Comparing the ratio of proboscis contacts on the pattern features to contacts on the pattern background during nectary search demonstrated that successful foraging bouts were associated with more precise pattern feature contacts (GLMM X ^2^ = 298.6, p = 0; Fig. 2b). The feature inspection ratios were generally higher for the di-and monosymmetric patterns on the first day (Fig. 2c). However, this trend was not significant, and pattern type was not found to significantly influence the feature inspection ratio. Instead, we found that the first pattern with which the moths interacted influenced their nectary search (GLMM first pattern type: X ^2^ =10.29, p=0.006, Fig. 2d), with inspections on the monosymmetric pattern on the first day leading to significantly more precise pattern feature contacts throughout all trials compared to interacting with the polysymmetric pattern.

To assess feeding efficiency across pattern types, we compared time to feeding, proboscis path tortuosity, and proboscis speed in the bouts with successful feeding. The interactions between the pattern type and day did not significantly influence these measures, so bouts across all days were compared between pattern types. The pattern inspection duration up to feeding in successful bouts was influenced by the pattern type (GLMM pattern type: X ^2^ =6.07, p=0.007), with significantly lower inspection times for the monosymmetric pattern with respect to the polysymmetric pattern (Fig. 3e, Table S). The order of pattern presentation was also an influencing factor in the inspection times (GLMM first pattern: X ^2^ =14.01, p<0.001), with moths taking longer to feed when presented with the dissymmetric pattern on the first trial day, compared to when they were presented with the polysymmetric pattern on the first day (Fig. S3a). The time to feeding was also influenced by the trial day (GLMM trial day: X ^2^ =6.36, p=0.041), with the moths on the third day taking significantly longer to find the nectary compared to the first day (Fig. S3a).

The inspection path tortuosity up to feeding was similarly influenced by pattern type (GLMM pattern type: X_2_^2^ =7.67, p=0.021), with significantly more tortuous paths on the polysymmetric pattern compared to the monosymmetric pattern (Fig. 3f). First pattern presented (GLMM first pattern: X_2_^2^ =10.14, p=0.006) and day (GLMM trial day: X_2_^2^ =8.6, p=0.013) also influenced the inspection path tortuosity. Path tortuosity was greater on the third trial day than on the second trial day (Fig. S3b), and experiencing the dissymmetric pattern first resulted in paths with higher tortuosity than being first presented with the monosymmetric pattern (Fig. S3b).

Pattern type also influenced the median proboscis speed (X ^2^ =16.27, p-value < 0.001), with significantly higher speeds on the dissymmetric pattern compared to both the mono-and polysymmetric pattern (Fig. 3g). Speed also varied with the trial day (X ^2^ =11.81, p-value = 0.002), with significantly higher inspection speeds on the third trial day compared to the second and first day (Fig. S3c).

Together, the results show that the innately preferred monosymmetric pattern in the horizontal plane benefited foraging efficiency and overall success. The preferred pattern guided the proboscis more directly to the central nectary, contributing to foraging efficiency. Interactions with the monosymmetric flowers also improved the hawkmoths’ pattern inspection precision in subsequent foraging bouts, suggesting a role for experience in moderating flower inspection.

We also assessed the effect of pattern symmetry on floral approach before the moths began inspecting the surface for the nectary. The approach direction was assessed by calculating the heading angles of the moths during flight. The median heading angles with respect to the pattern centre during approach were concentrated along the same angles as the outer pattern arm features (Fig. S3d), however, the angles were not significantly heterogeneously distributed nor different between pattern types. The distribution of the first proboscis contacts was found to be significantly heterogeneously distributed in all except the polysymmetric pattern. Significant differences in the distribution of contact angles were also seen between the polysymmetric pattern and both, the di-and monosymmetric patterns (Fig. 3h). There were no significant differences between the dissymmetric and monosymmetric patterns. Additionally, in the monosymmetric pattern, the arms on the central axis (Fig. S3e) received significantly more contacts than the sideward arms, aligning the contacts predominantly along the central axis of the pattern. A significant difference between proboscis contacts was also found on the arms of the polysymmetric pattern, but since each arm in this pattern presents the same visual information, this could not have been caused by a selective preference for a particular feature. Rather, this may have been an artefact of the moths preferring a particular location in the cage to approach the patterns from. The pattern features thus aligned the moths’ position across a single axis in the di-and monosymmetric patterns, and across two axes in the polysymmetric pattern.

Thus, the naively preferred symmetry types in the horizontal display plane guided floral inspection to achieve greater foraging success. The di-and monosymmetric patterns also guided the approach flight and the initial pattern contact to receive more directionally consistent inspections.

### Foraging on vertically displayed bar orientation patterns

The influence of the naively preferred patterns in the vertical display plane on foraging behaviour was assessed by tracking moths as they foraged on a single vertically presented pattern. Since feature position and feature orientation on the patterns emerged as the relevant factors influencing floral pattern choice (Fig. 1e,f), approach flight and flower inspection were compared across the 0° and 90° bars from the bar orientation set, and across all patterns in the feature position pattern set.

The foraging interactions with the 0° and 90° bars were filmed from the side to assess movement in the vertical plane, and from the top to assess movement in the lateral plane during foraging (Fig. 3a). The approach trajectories across both patterns revealed similarly stereotyped flight trajectories in the vertical plane (Fig. 3b). In the flight cage, moths approached the pattern by descending downwards from the top of the cage towards the central region, at which point the first proboscis contact with the pattern occurred. Upon contact, the hawkmoths moved upwards. This was reflected in the difference in displacement in the vertical plane before and after the first contact (LMM displacement instance: X ^2^ =152.60, p=0; Fig. 3d), with downward displacement or hovering 0.5s before the initial proboscis contact and upward movement 0.5s after the initial contact. In the lateral plane, the approach trajectories were much more variable and did not reveal a clear stereotypy (Fig. 3c). Across pattern types, there was a significant difference in displacement with relatively more leftward movement during approach and rightward movement during initial inspection (Fig. 3d). Since both patterns are symmetrical around the vertical plane, this might have been caused by some spatial bias in the flight cage, leading animals to preferentially approach from one side. Between the bar orientation types, there were no significant differences in displacement 0.5 s before first contact in either plane, but the patterns significantly influenced displacement 0.5 s after the first contact (Fig. 3d). In the vertical plane, the moths covered greater upward distances during inspection of 90° bar orientations (t (155): t=-1.08, p=0.004). While there was no difference in mean displacement in the lateral plane between bar orientations, the distribution of inspection distances differed between the pattern types (Levene’s test: F (1,78), F=13.30, p=0.0005), with greater variability in the 0° bar orientation pattern.

We assessed the approach trajectory dynamics to check whether pattern orientation type influenced the flight path and stability. Approach tortuosity in the vertical plane was similar across pattern types, but in the lateral plane the approaches to the 0° bar orientation pattern were significantly more tortuous (Wilcoxon’s test: W= 793, p = 0.0004; Fig. 3e). To quantify flight path jitter, we assessed the frequency distribution of the Fourier-transformed approach paths in both view planes (Fig. S4e,g). A majority (99%) of the trajectory components from both views were composed of frequencies below the 5 Hz range and were not significantly different between the pattern types in either view plane (Fig. S5f,h).

To quantify whether nectary inspection was influenced by the bar orientation type, we assessed the position of the proboscis at first contact. Following the differences in approach trajectories across pattern types, the first proboscis contacts were distributed differently between the patterns in both the vertical (Levene’s test: F(1,83), F=50.95, p=0) and lateral plane (Levene’s test: F(1,85), F=71.18, p=0; Fig. 3f). First contacts occurred across the whole bar feature in both patterns. However, while they occurred on the bar for the 90° orientation pattern, they were frequently positioned below the bar for the 0° orientation pattern (Fig. 3f, S4 a,b). To quantify whether a nectary contact occurred within a foraging bout, the movement of the proboscis through the pattern centre was assessed. The total proportion of bouts in which a central contact occurred did not differ significantly between the pattern types (Fig.3g). While centre contacts occurred slightly earlier with the 90° bar orientation pattern, this difference was not significant (Fig.3g).

Thus, we show that the hawkmoths had a stereotyped foraging movement in the vertical plane but not in the lateral plane for both pattern types. The bar orientation type guided proboscis and body movement during pattern inspection. However, we did not find significantly improved foraging success with the preferred 0° bar orientation pattern.

### Foraging on vertically displayed feature position patterns

In the feature position pattern set, the foraging interactions were filmed from the side to record the vertical movement and compare the stereotyped approach flights and interactions with the patterns (Fig. 4a). Similar to the flight trajectories in the orientation pattern set, the direction of displacement was significantly different before and after contact across all pattern types (LMM displacement instance: X ^2^ =149.84, p=0; Fig. 4b). Displacement did not significantly differ across pattern types. The approach path tortuosities were also not significantly different across pattern types (Fig. 4c).

The pattern type, however, influenced the location of the first proboscis contact on the pattern (GLMM pattern: X ^2^ =77.94, p=0) with contacts significantly in the lower half of the disc for the patterns with higher contrast elements in the lower half (Fig. 4d). Pattern type also significantly influenced the number of bouts in which the centre was contacted (X ^2^ =9.04, p=0.03) with more successful bouts on patterns with additional features on the lower half (Fig. 4e), although pairwise differences were not significant. The higher proportion of centre contacts in these patterns resulted from the lower position of the body and proboscis at first contact (Fig. 4d, S5a) and the subsequent stereotyped upward direction in which the moth moved to inspect the pattern.

Thus, across vertically displayed patterns with different feature positions, the naively preferred pattern with greater contrasting features in the lower half shifted the stereotyped approach trajectories so the first proboscis contact occurred at or below the central pattern region. Subsequently, during the upward inspection, there was a greater likelihood of contacting the central nectary.

### Characteristics of hawkmoth-foraged flowers

To compare the foraging choices of hummingbird hawkmoths on artificial flowers with the display planes and floral symmetry of plant species visited by these hawkmoths in nature, we analysed flower visitation data based on citizen science reports (see Methods). In addition to flower symmetry and display, the position of patterns on the flowers was annotated to compare with feature preferences on the artificial flowers. Hummingbird hawkmoths were recorded visiting 170 plant species across 50 families (Fig.5 b,c) in Germany. The highest number of plant species visited was recorded in the Lamiaceae family (20%), followed by plants in the Asteraceae family (10.6%). Across these species, radial (49.4%) and bilateral (38.8%) flower symmetries were represented, along with filled flowers (11.8%) with no distinct plane of symmetry (Fig. 5b). Floral displays ranged from primarily vertical (61.8%), primarily horizontal (22.9%), downward-facing (2.9%), and mixed planes of display (12.4%) (Fig. 5c). Patterns were displayed by 57.06% of the recorded flower species.

In order to compare flower features with the patterns and displays used in the naïve choice assays, plant species with flowers that were classified as primarily vertical or primarily horizontal (based on the display planes of over 70% of flowers on a plant), and symmetries that could be grouped as either radial or bilateral, were quantified in the subsequent analysis. Within this set, horizontally displayed flowers were found to most frequently display radial symmetry (64.1%), while vertically displayed flowers had a greater proportion of species with bilaterally symmetric flowers (60%) (Fig. 5d). Interestingly, this was contrary to the hawkmoths’ naïve symmetry preferences with horizontally displayed flowers, where they showed a greater preference for bilaterally symmetric patterns (Fig. 1b).

Within the bilaterally symmetric flowers that were vertically displayed, 61.9% displayed more visually prominent ventral petals known as lower lips (Fig. 5e). Amongst vertically displayed bilateral flowers with patterns, the patterns were located either around the flower face (52.4%) or in the bottom half (40.5%), but rarely on the top half (Fig. 5f). Such flowers with prominent visual contrast on the ventral half were comparable to the naively preferred artificial patterns with features on the lower half (Fig. 1f)

From assessing natural flower visitation, hawkmoths were found to be generalists that visit flowers with various visual traits. Amongst the visited flower species, horizontally displayed flowers were largely radially symmetric, while vertically displayed flowers tended to have bilateral symmetry. The majority of vertically displayed bilateral flowers displayed more prominent visual features in the ventral half of the corolla, with patterns, a large lower lip, or both. In comparison with the naïve pattern choices, the dataset contained several examples of flowers that resembled the preferred vertically displayed patterns with ventral visual contrast. However, the preferred bilateral symmetry in the horizontal display was not strongly represented in the visited flowers.

## Discussion

In this study, we tested naïve hawkmoths for symmetry preferences during flower choice and found that pattern preferences were linked to the display plane of the patterns. While symmetry was relevant for pattern selection in the horizontal display, the orientation and position of the pattern features, rather than symmetry type, guided choice in the vertical display plane. Assessing the functional relevance of these naïve preferences in relation to foraging revealed that the preferred pattern features enabled more efficient foraging. The horizontally displayed di-and monosymmetric patterns guided proboscis inspections to the food more efficiently, while the preferred features in the vertical plane influenced the initial proboscis contacts to create a higher likelihood of interacting with the central nectary. Flowers naturally foraged on by hawkmoths did not entirely replicate their innately preferred patterns-horizontally displayed flowers were typically radial, i.e. polysymmetric, unlike the monosymmetric artificial flowers that were preferred by the naïve moths. However, several plant species with flowers in the vertical plane had similar features as the preferred vertical artificial patterns, with visually prominent lower halves.

### Symmetry preferences for flower choice

Hummingbird hawkmoths were found to discriminate between symmetry types during flower choice with no prior foraging experience in our study. Previously, flower foraging insects such as honeybees (Giurfa et al., 1996) and bumblebees (Plowright et al., 2011) have been shown to learn symmetry through rewarded associations. In hoverflies (Mishra et al., 2025; Møller & Sorci, 1998), honeybees (Giurfa et al., 1996; Wignall et al., 2006) and bumblebees (Møller, 1995; Rodríguez et al., 2004; however, see Plowright et al., 2011) unlearned preferences for symmetric over asymmetric foraging targets have also been described. However, evidence for foraging preferences across symmetry types, ie, radial and bilateral, is scarce, with one study where honeybees were shown to select radially symmetric flowers over transversely bilateral flowers (Wignall et al., 2006). In the present study, we show that the hawkmoths naïvely preferred a bilateral pattern with a single plane of symmetry over a radial pattern. With insect orders across Hymenoptera, Diptera, and now Lepidoptera found to discern symmetry as a visual feature, symmetry is likely to be ubiquitously perceived by insects that forage on flowers. Differences in floral symmetry preferences between insects with disparate foraging behaviours could indicate pollinator specialisation resulting from conserved foraging interactions within specific plant and pollinator species.

Interestingly, the bilateral pattern preferences recorded in the horizontally displayed artificial flowers were not replicated when the same flowers were presented vertically. In selecting vertically displayed patterns, the hawkmoths preferred features on the lower half of the patterns and also assessed the local feature orientations within the pattern. The differences in pattern preferences between the horizontal and vertically displayed artificial flowers could have arisen due to the differences in the images being viewed across the display types. One such difference would arise from the rotationally fixed viewpoint in the vertical display, whereas in the horizontal display the hawkmoth could view the pattern in various rotated forms. However, the whole symmetry or global spatial arrangement of the pattern had to be assessed in the horizontal plane to preferentially select the monosymmetric flower pattern. Since the same symmetric properties were conserved in the vertical plane, differences in pattern choices between the vertical and horizontal displays cannot be explained by rotationally different patterns being viewed across display planes. For the same reason, image differences arising from shear or distortion effects across the display planes also would not explain the pattern choices. Another outcome of varying the display planes would be a change in the region of the eye upon which the images are subtended, which would primarily be on the ventral eye region for the horizontally displayed patterns and primarily on the frontal eye region for the vertically displayed patterns. In honeybees, different areas of the visual field have been found to differ in whether they could learn patterns, with orientation being learnt in the frontal and lateral region but not the dorsal or ventral region (Giger & Srinivasan, 1997). Differences in stimulus display were also shown to influence the speed and accuracy of learning colour and orientation discrimination in this species (Wolf et al., 2015). Smaller differences in visual cue reliance across the visual field during flight have also been described in some moth species (Copley et al., 2018; Vickers & Baker, 1994) as well as in the fruit fly (Mazo & Theobald, 2014) but uniquely, in hummingbird hawkmoths, visual motion cues in different parts of the visual field were shown to produce considerably different responses, from flight stabilisation to eliciting avoidance behaviour (Bigge et al., 2021, 2025). Such widespread occurrences of varied responses to visual stimulation in different parts of the insect eye could suggest that visual information is perceived or ‘seen’ differently in different parts of the eye. Alternatively, the same cues could be perceived but may have different valences across the eye. The latter explanation, in the present case, would suggest that symmetry was similarly detected in both display planes, but orientation and feature position-pattern qualities that remain constant only in the vertical axis-held greater relevance for flower choice in the vertical display.

A comparative analysis of display-dependent visual processing across insect species may reveal further instances of such asymmetries in choice, which would be an essential factor to consider when interpreting insect sensory behaviour.

### Foraging function of innate pattern preferences

The differences in innate preferences in the hawkmoths were associated with the distinct foraging benefits afforded by the preferred flower patterns in their respective display planes. These varied foraging benefits were related to how the hawkmoths’ approach and inspection were influenced by the floral display. During approach, movement towards the vertically displayed flowers was strongly stereotyped, whereas no such stereotypy was observed for the close-range approaches to horizontally displayed flowers, where the forager could approach from any direction. For foraging-naïve hawkmoths, the preferred bilaterally symmetric patterns were found to guide more direct paths to the nectary when horizontally presented, potentially by constraining the nectary search area across a single axis instead of maintaining multiple possible inspection regions. The increased foraging success afforded by the preferred bilateral patterns, along with the small but significant improvement in foraging efficiency, could have led to the better foraging performance of these individuals across subsequent trials. However, for flowers displayed vertically, the motor guidance associated with the monosymmetric pattern would not be advantageous since the hawkmoths’ approaches were already restricted across the flower’s dorsoventral axis. Similar effects on foraging approach have also been described in bumblebees, where the vertical display, rather than stimulus shape, strongly constricted the forager’s approach angles (Jirgal & Ohashi, 2023) and where bilateral shapes in the horizontal display slightly reduced approach variability (Culbert & Forrest, 2016).

Besides approach, the floral display type also influenced the motor control of the proboscis during flower inspection. The proboscis contact on horizontally displayed flowers was initiated on the outer edge of the flower, following which the proboscis tip moved inwards toward the centre (Fig. S7). In vertically displayed flowers, on the other hand, the initial proboscis contact occurred at or below the centre of the flower rather than at the floral edge, after which the proboscis inspection was directed upwards towards the flower’s dorsal edge. Such differences have been described in a hoverfly species, where after landing, movement is directed around the centre for horizontally displayed flowers and across the flower when it is vertically displayed (Wiegel & Lunau, 2026). For hawkmoths, the patterns that attracted significantly greater innate choice in the vertical display were the ones that split the flower into dorsoventral halves, such as the 0° oriented bar pattern. The dorsoventral split of the flower guided the initial proboscis contact below the centre of the flower, which improved the individual’s probability of contacting the central nectary while moving upwards. This advantage of initiating proboscis inspection below the centre of the flower was also afforded by the patterns with greater feature contrast in the lower half. The patterns tested with differential dorsoventral contrast features additionally offered a pattern centre that increased the probability of contacting the centre overall compared to the bar patterns, suggesting an additional benefit to foraging from flower patterns that mark the floral centre.

Together, the association of the patterns with foraging benefits suggests a strong interaction between visual preferences and motor behaviours during foraging. Selection on innate pattern preferences was likely shaped by the kinematics of hawkmoth flight and probing and, in turn, visual preferences for rewarding floral features may have produced stereotyped foraging movements.

### Linking foraging behaviours and flower features

Flowers that were naturally foraged on by hummingbird hawkmoths were found to share some visual characteristics with the hawkmoths’ innately preferred patterns. Specifically, vertically displayed bilateral flowers had prominent visual features in the ventral half in the form of contrasting patterns, larger petals, or both, which likened the innately preferred patterns with prominent ventral features. Interestingly, flower characteristics such as the larger ventral petal, also known as the labellum or ‘lower-lip’, have been described in flowers foraged by hummingbirds (Smith et al., 1996; Temeles & Rankin, 2000) and bumblebees (Castellanos et al., 2004; Zung et al., 2015) where these petals were found to increase foraging success.

The effect of bilaterally symmetric flowers with lower-lip petals effectively guiding the pollinator approach could represent the pollen-placement hypothesis (Dafni & Kevan, 1996; Fenster et al., 2004)-where plants invest in depositing concentrated pollen loads in precise locations, and the floral traits of such plants shape the foragers’ approach and interaction to receive reliable pollination. Specialisation of pollen placement has specifically been associated with vertically oriented bilateral flowers, which are found to guide the forager’s approach and interaction (Fenster et al., 2009; Jirgal & Ohashi, 2023; Wiegel & Lunau, 2026). A greater salience for features in the ventral half of patterns has also been previously reported from visual learning experiments in honeybees and bumblebees (Giurfa et al., 1999; Guiraud et al., 2018; MaBouDi et al., 2023). In these studies, the bees memorised the ventral features rather than the whole pattern, and visual scanning behaviours were also predominantly in the ventral section of the patterns. The stereotyped inspection of ventral floral features among insect foragers such as hawkmoths and bees could have exerted selective pressure on host plants to place the pollen above the nectary, so pollen is deposited on the back or head of the forager. This is the case for plant families like Lamiaceae, which have nototribic or dorsal pollen deposition (Claßen-Bockhoff, 2007), a group that was majorly represented in the dataset of hawkmoth-foraged flowers.

The inspection paths taken by the hawkmoths from the lower to the upper half of the pattern, in addition to enabling nectary contacts, could also be a good foraging strategy for clustered inflorescences where resource allocation typically decreases across the flowers from the proximal to the distal ends (Diggle, 1997; Kliber & Eckert, 2004). Such upward foraging inspections on vertical inflorescences have been described in bumblebees and wasps (Corbet et al., 1981; Pyke, 1978; Waddington & Heinrich, 1979) and in a species of hoverfly (Wiegel & Lunau, 2026). The exception to the upward foraging movement was described in bumblebees that engaged in nectar robbing of flowers (Corbet et al., 1981). Thus, features that guide upward movement may be advantageous to both pollinators and plants.

A contradicting result between innate pattern preferences and the flower dataset was that only a single plant among over 100 species represented the horizontally displayed bilateral symmetry that attracted the most innate foraging choices in the hawkmoths. The species, *Valeriana rubra*, has clustered inflorescences of pink or white flowers, originates in the Mediterranean region, and has been introduced as a common garden species in northern Europe. The disparity between the innate choice and floral feature data, in this case, could be due to the regional mismatch of annotating floral data in Germany if hummingbird hawkmoth pattern vision coevolved with plants in the Mediterranean region. Hummingbird hawkmoths are a seasonally migratory species that fly to northwest Europe in spring and overwinter in the Mediterranean region. With their oviposition host plants, the Galium sp., distributed across their migratory region (Ortega-Olivencia & Devesa, 2003), their life cycle is not geographically restricted. However, the warmer months are more conducive to reproduction, with faster developmental periods (Pfaff & Kelber, 2005) and more foraging resources, which makes it likely that new generations emerge in northwestern Europe, from where the flower foraging data was annotated. Thus, a regional mismatch of data is unlikely. However, the distribution of plant species in Northern Europe has greatly changed over the last century with human-driven horticultural practices such that annotated flowers may also reflect flowers learned through opportunistic foraging rather than being representative of innate preferences. Additionally, while the present analysis quantified the range of visual features visited by the hawkmoths, the visitation abundance to individual plant species, which are better indicators of preference, could not be quantified. The mismatches found in the floral dataset could also be an important indication that innate foraging preferences may not necessarily develop to represent foraging targets and enable flower recognition but instead could have emerged to aid efficient movement in naïve foragers. In nature, deviations in floral radial symmetry might be used by naïve foragers to guide their initial foraging to locate the nectary, after which they could transfer these learnt associations while foraging on a diverse range of flowers with centrally placed nectaries.

## Conclusions

In hummingbird hawkmoths, visual patterns drove innate flower choice in addition to serving as both approach and nectar guides. The correspondence between the hawkmoths’ visuomotor foraging behaviours and foraging success indicates that their sensory biases were shaped by plant-pollinator interactions. The selection that drove such visuomotor biases was likely mediated by both plant and pollinator, since these biases could also benefit pollination success in the host plants. The parallels in foraging movement that could be drawn between the hawkmoths and other nectivorous insects suggest a convergence in foraging behaviours in response to flower display rather than representing traits unique to the hover-feeding behaviour of the hawkmoths.

Thus, in this study, we highlight the role of floral patterns for foraging interactions and provide evidence that innate sensory biases are shaped by ecological function.

## Author contributions

Conceptualization: A.N.M., A.S.; Methodology: A.N.M., A.S., L.W. Validation: A.N.M., A.S.; Formal analysis: A.N.M; Investigation: A.N.M., A.S.; Resources: A.S.; Data curation: A.N.M.; Writing - original draft: A.N.M.; Writing - review & editing: A.N.M., A.S.; Visualization: A.N.M.; Supervision: A.S.; Funding acquisition: A.S

## Acknowledgements

We are grateful to Alexander Dietz and NABU, Naturgucker (https://nabu-naturgucker.de/), together with the people who contributed their observations, for providing the floral dataset in this study. We thank Yvonne Hertenberger for assistance with collecting data, and all the animal caretakers who helped maintain the hawkmoth populations.

