## Supplementary Information for "A hawkmoth’s point of view - Innate symmetry preferences are informed by floral display and foraging function in an insect pollinator"

##### Figures

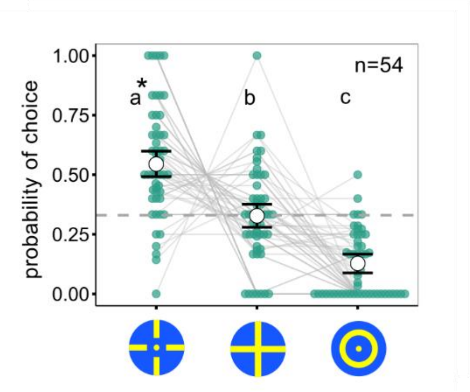


Fig. S1 Probability of choice across a full and disrupted cross, and a concentric circle pattern. Error bars indicate 95% confidence intervals. The number of hawkmoths tested in each pattern set is given by 'n'. Letters represent significant differences between pattern types, and the patterns chosen significantly above random chance are denoted by asterisks (*$p < 0.05$).

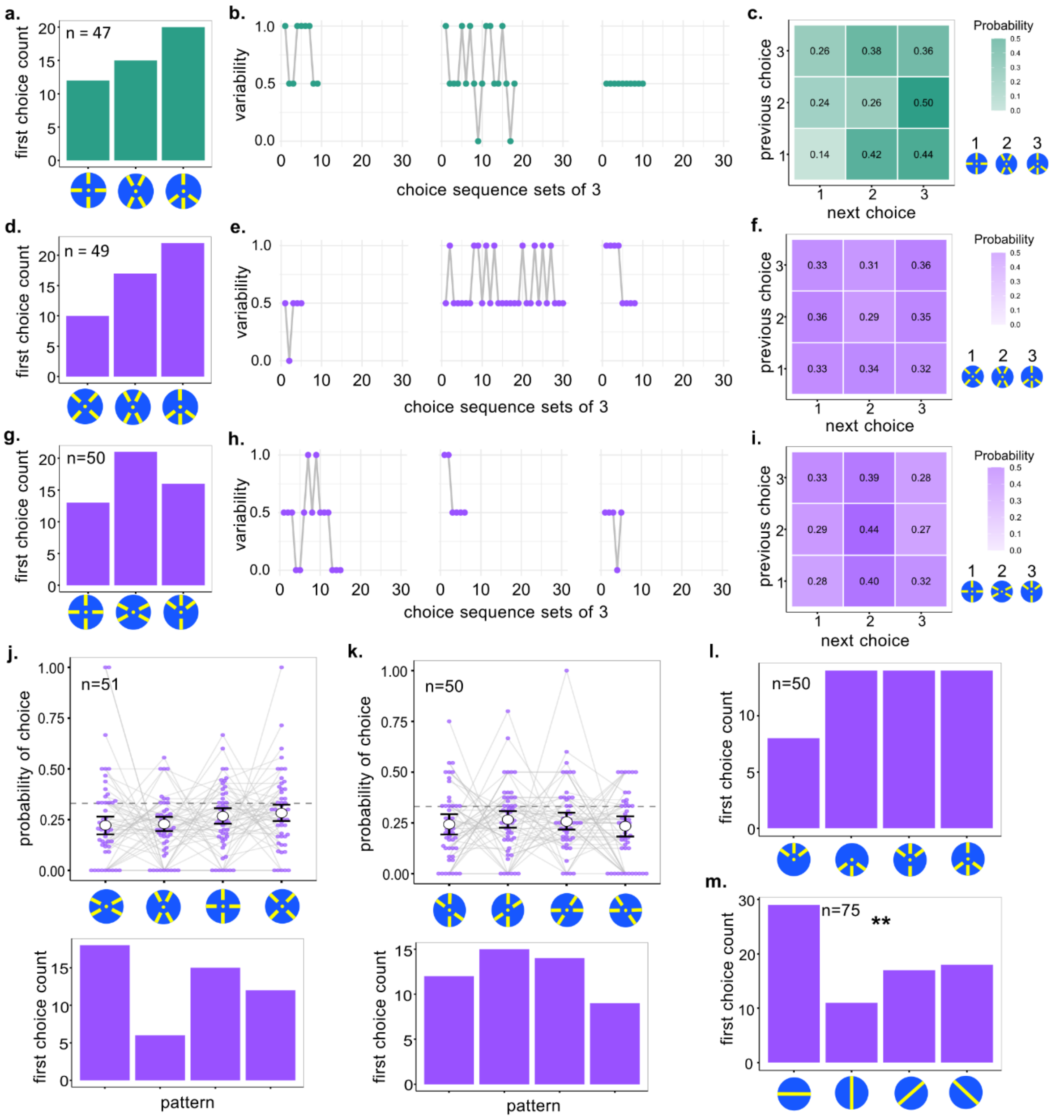


Fig. S2 **a, d, e**. First choices across symmetry patterns in the horizontal and vertical display planes. **b, e, h.** Variability in pattern choices calculated across a rolling window of 3 choices throughout the choice sequence, where 0 indicates choosing the same pattern thrice in a row, and 1 indicates that three different patterns were sequentially chosen. Choice variability is visualised for three individuals in each set of symmetry patterns in the horizontal and vertical display planes. **c, f, i.** Transition matrices denoting the probability of contacting a specific pattern based on the previous choice. **j, k.** Probability of naïve pattern choices by hawkmoths across poly- and dissymmetric patterns at two different orientations (j.) and across dissymmetric patterns at four different orientations (top). Posterior predicted means are represented by white circles with error bars representing 95% Credible Intervals. Coloured points represent proportion of choice from individual hawkmoths, with the grey lines connecting choice proportions across the same individual. Distribution of first choices for the different pattern sets are shown at the bottom of the respective patterns. **l.** Distribution of first choices across the feature position pattern set. **m.** Distribution of first choices across the bar orientation pattern set. Asterisks denote results of a Chi-square test (p<0.01).

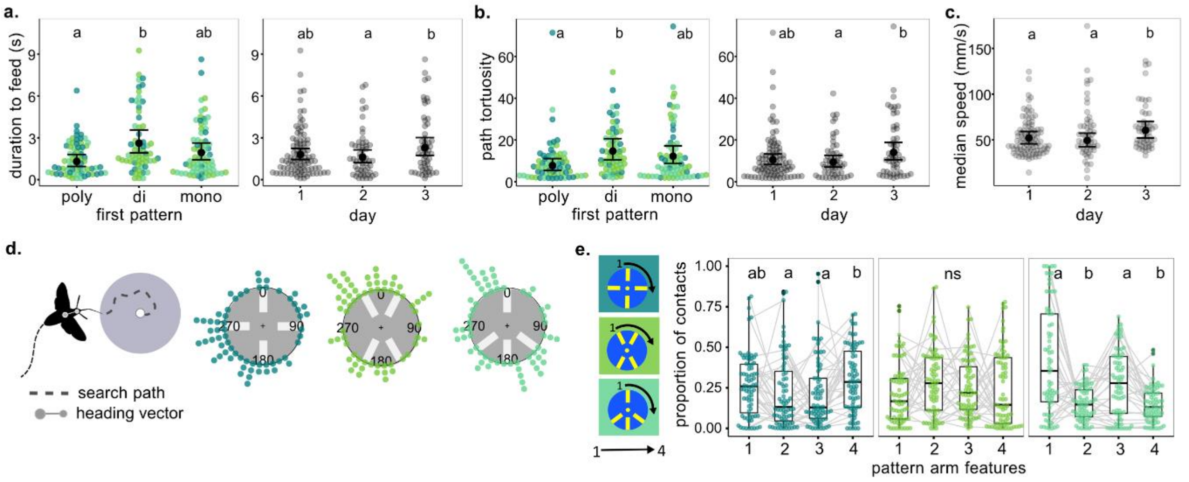


Fig S3 **a-c.** Proboscis path dynamics across symmetry types from bouts with feeding. Distribution of time taken to feed from the central nectary (a.)**,** tortuosity of the proboscis path (b.), and mean proboscis speed (c.) with predicted means from the GLMM denoted by white circle and the error bars depicting 95% Confidence Intervals. Plots on the left show data grouped according to symmetry type interacted with on the first day. Colours in the data points correspond to the symmetry type involved in the bout. Pattern types with no differences (p> 0.05) share the same letter. **d.** Distribution of median heading directions during approach from a close range (60 mm from pattern centre) with respect to the centre of the pattern. Proportion of inspection bouts on the first trial day with successful feeding. *e.* Ratio of proboscis contacts within each yellow arm of the pattern across symmetry types. The arms denoted by the respective numbers are illustrated by the black arrow around each pattern type in the graphic to the left.

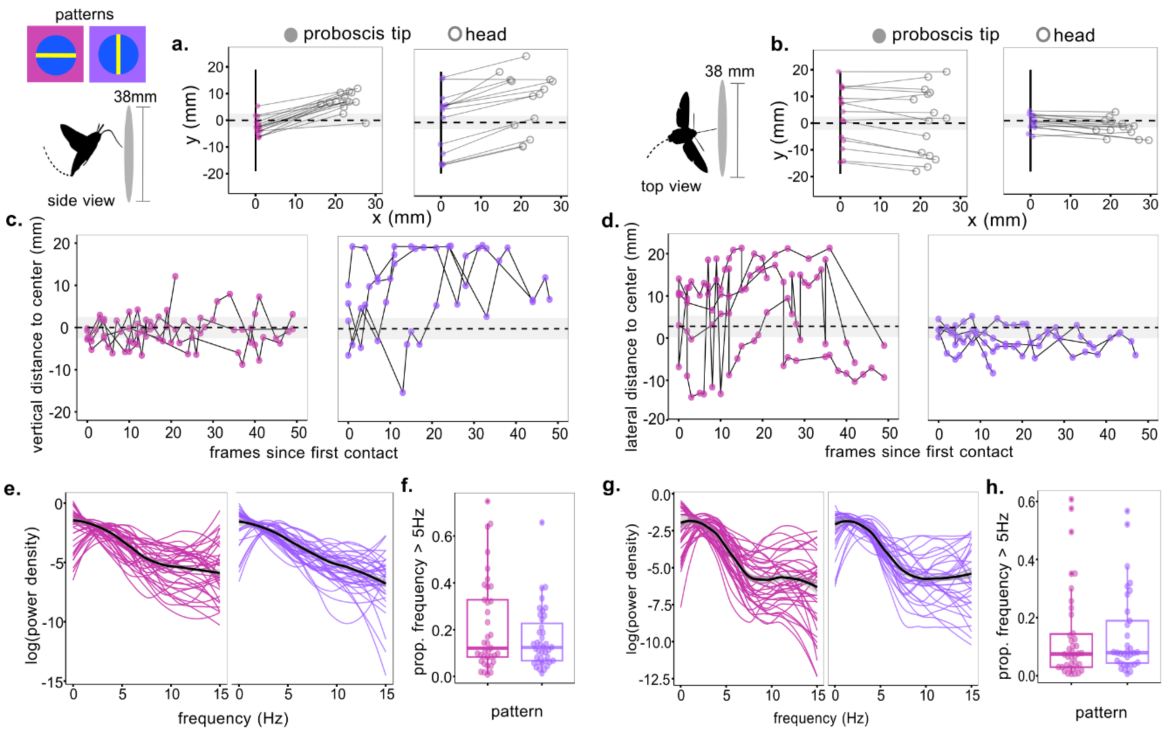


Fig. S4 **a, b.** Position of the hawkmoth with respect to the pattern (black vertical line) at first contact in the vertical plane (a.) and lateral plane (b) across the bar patterns. **c,d.** Proboscis contacts on the pattern across 50 frames after first contact. Proboscis movement is visualised for a single individual across different patterns in the bar pattern set. Contacts from the same bout are connected by a black line which is interpolated across frames where proboscis contacts were not annotated. **e, g.** Power density of Fourier-transformed approach trajectories to the bar orientation patterns in the vertical (d.) and lateral (f.) plane. Coloured lines indicate individual trajectories and the black line gives the median power densities. **f, h.** Proportion of frequencies greater than 5 Hz (jittered paths) from the Fourier-transformed approach trajectories across the pattern types.

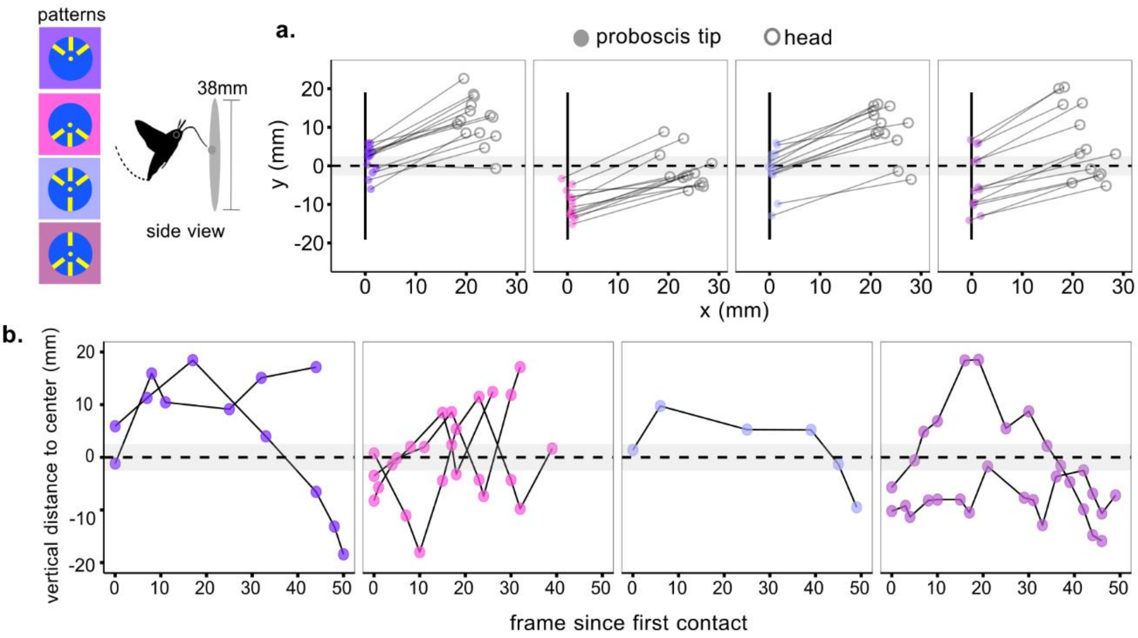


Fig. S5 **a**. Position of the hawkmoth with respect to the pattern (black vertical line) at first contact in the vertical plane across the feature-position patterns. **b**. Proboscis contacts on the pattern across 50 frames after first contact. Proboscis movement is visualised for a single individual across different patterns in a pattern set. Contacts from the same interaction bout are connected by a black line, which is interpolated across frames where proboscis contacts were not annotated.

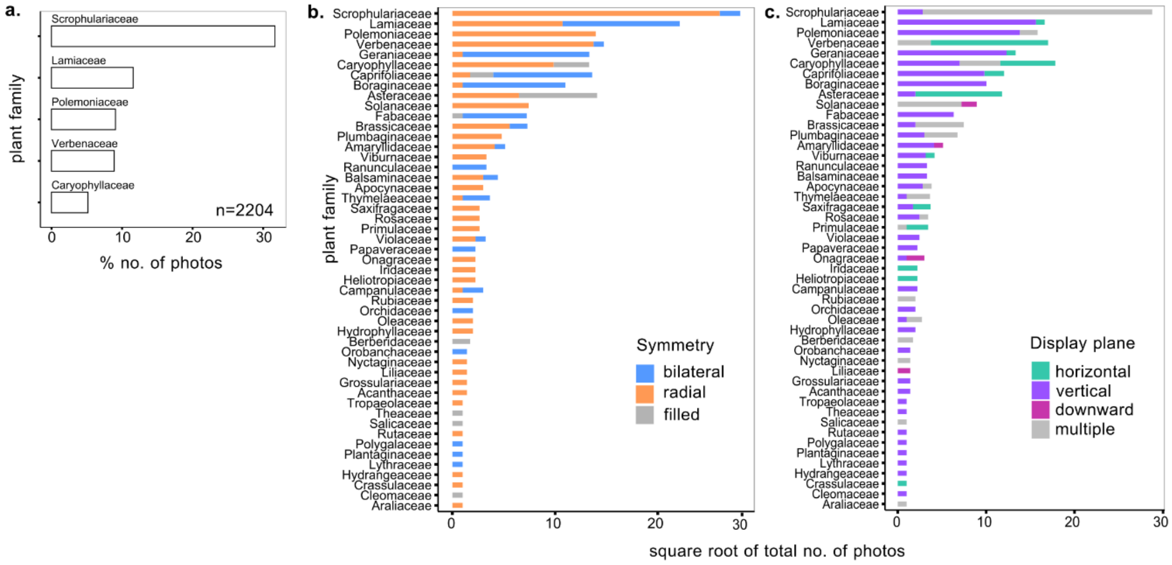


Fig. S6 **a**. The proportion of photographs across plant families containing over 0.05% of the analysed dataset. 'n' represents the total dataset of photographs with flower-hummingbird hawkmoth interactions. **b, c**. Number of photos with foraging visits across plant families. Colours depict symmetry type (b.) and display plane (c.) of the flowers.

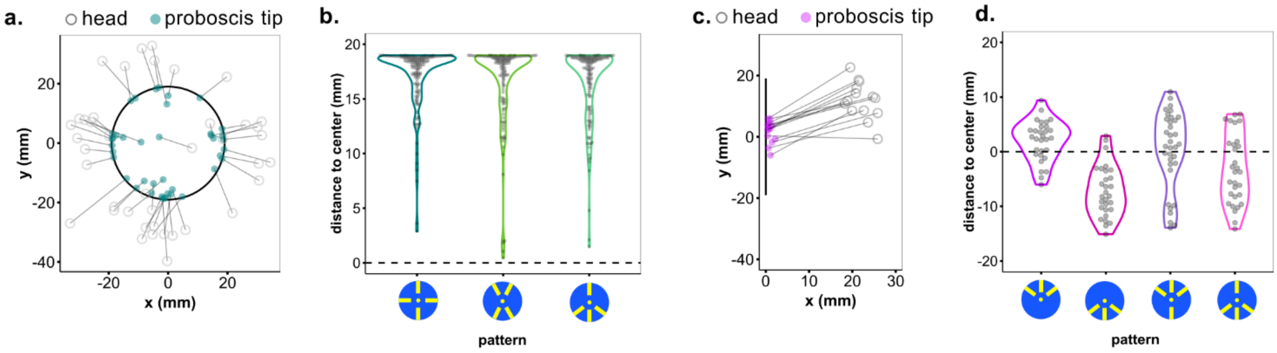


Fig. S7 **a.** Proboscis tip and head coordinates of the hawkmoths around the stimulus border for the horizontally displayed patterns during the first contacts. **b.** Distance of the proboscis tip from the centre of the patterns at first contact on the horizontally displayed patterns. **c.** Proboscis tip and head coordinates of the hawkmoths from the side view for the vertically displayed patterns during the first contacts. **d**. Distance of the proboscis tip from the centre of the patterns at first contact on the vertically displayed patterns. Positive values depict the contacts above the pattern centre and negative values depict contacts below the pattern centre

### Tables

*Table S1. Estimated probability of naïve pattern choice within a pattern set along with the adjusted p-values of the patterns being chosen above random chance. Significant p-values are highlighted in bold.*

| **Display position** | **Pattern set** | **Pattern type** | **Odds ratio** | **Estimate [95% Credible Interval]** | **P value** |
|---|---|---|---|---|---|
| horizontal | symmetry | polysymmetric | 0.30 | 0.23 [0.19, 0.26] | 1 |
| | | dissymmetric | 0.53 | 0.34 [0.31, 0.38] | 0.41 |
| | | monosymmetric | 0.74 | 0.43 [0.39, 0.47] | **0** |
| vertical | symmetry set 1 | polysymmetric | 0.49 | 0.33 [0.28, 0.38] | 0.81 |
| | | dissymmetric | 0.46 | 0.31 [0.27, 0.36] | 0.81 |
| | | monosymmetric | 0.56 | 0.36 [0.31, 0.41] | 0.39 |
| | symmetry set 2 | polysymmetric | 0.40 | 0.29 [0.24, 0.34] | 0.96 |
| | | dissymmetric | 0.73 | 0.42 [0.37, 0.48] | **0.002** |
| | | monosymmetric | 0.41 | 0.29 [0.24, 0.34] | 0.96 |

| | | | | | |
|---|---|---|---|---|---|
| | bar orientation | 0° | 0.49 | 0.33 [0.26, 0.39] | **0.03** |
| | | 90° | 0.27 | 0.21 [0.16, 0.27] | 0.98 |
| | | 45° | 0.36 | 0.27 [0.20, 0.34] | 0.64 |
| | | -45° | 0.24 | 0.19 [0.14, 0.26] | 0.98 |
| | feature position | only top | 0.20 | 0.17 [0.13, 0.21] | 1 |
| | | only bottom | 0.30 | 0.23 [0.18, 0.28] | 1 |
| | | greater top | 0.40 | 0.28 [0.23, 0.33] | 0.27 |
| | | greater bottom | 0.47 | 0.32 [0.26, 0.38] | **0.03** |
